# Disease associated human TCR characterization by deep-learning framework TCR-DeepInsight

**DOI:** 10.1101/2023.05.22.541406

**Authors:** Ziwei Xue, Lize Wu, Ruonan Tian, Zuozhu Liu, Yadan Bai, Di Sun, Yixin Guo, Pengwei Chen, Yu Zhao, Bing He, Lie Wang, Jianhua Yao, Linrong Lu, Wanlu Liu

## Abstract

T cell function is defined by both T cell receptors (TCR) and T cell gene expression (GEX). Although single-cell technology enables the simultaneous capture of TCR and GEX information, the lack of a reference atlas and computational tools hinders our ability to uncover the fundamental TCR usage rules and to efficiently characterize disease-associated TCRs (dTCR). Here, through the collection of million-scale single-cell GEX-TCR reference atlas comprising 20 diverse disease conditions, we revealed the intrinsic features of TCR-MHC (Major Histocompatibility Complex) restriction in CD4/CD8 lineages. We observed the higher coherence for TCRα/β chains in memory T cells, and detected widely-existing public TCRα/β pairs across individuals. Building upon the reference atlas, we introduced TCR- DeepInsight, a deep-learning framework featuring a disease specificity scoring system that enables the characterization of dTCR clusters with similar GEX-TCR. Our study provides a valuable tool for researchers to analyze single-cell GEX-TCR data and identify dTCRs comprehensively and robustly.

## INTRODUCTION

T cells play a crucial role in the adaptive immune response by recognizing and eliminating pathogen-infected cells and providing cancer immunosurveillance. On the other hand, dysregulated T cell activation can also promote the pathogenesis of autoimmune diseases. In both protective immune and autoimmune responses, T cells exert their function in an antigen-specific manner, through the recognition of pMHC (peptide Major Histocompatibility Complex) via the T cell receptor (TCR) on the cell surface. The TCR consists of α and β chains with highly variable regions on both chains, contributing to the binding specificity between TCRs and antigen peptides. During their development in the thymus, T cells generate an enormous diversity of TCRs through V(D)J recombination and random nucleotide insertions and deletions in both α and β chains, enabling the recognition of an almost infinite number of pathogen-derived or self-antigens. Recent estimates of the total diversity of TCRs range from 10^20^ to 10^61^.^1,2^

In response to infections, cancer progression, or autoimmune diseases, T cells undergo clonal expansion, resulting in the production of a large number of T cells with identical TCRs that inherit the same antigen specificity. The characterization of TCR repertoire is critical for understanding the mechanisms of immune responses and developing effective immunotherapies. There are three complementarity-determining regions (CDRs) on both TCR α and β chain, with CDR3 regions directly binding to antigen peptides and thus exhibiting greater diversity compared to CDR1 or CDR2.^3^ Early spectratyping of TCRβ reveals preferential V and J segment usage and perturbations in CDR3β pattern in clonally expanded clonotypes in autoimmune diseases including Rheumatoid Arthritis (RA)^4^, Multiple Sclerosis (MS)^5^, and infectious diseases ^6,7^ However, these low-throughput and low-resolution analysis often biases the interpretation of Vβ and Jβ usage and CDR3β features. Next-generation sequencing (NGS) has revolutionized TCR repertoire analysis and led to a more concise understanding of TCRs in different disease conditions. ^8–11^ However, major drawbacks of NGS lie in the lack of TCRα/β chain pairing information and the incomplete coverage of TCR sequences. Single-cell immune profiling overcomes these limitations and has emerged as a powerful tool for analyzing paired TCRα/β repertoires at a single-cell resolution. Moreover, this technology captures the corresponding gene expression (GEX) profiles, allowing the linking of individual T cell clonotypes to their transcriptome dynamics during immune responses. Example studies using single-cell immune profiling include the identification of T cell clones associated with response to checkpoint inhibitors in patients with basal or squamous cell carcinoma before and after anti-PD-1 therapy^12^, the detection of T cell clonal expansion in COVID-19 infection^12–17^, and the analysis of TCR repertoire diversity in patients with autoimmune diseases.^18,19^ Sequencing-based technologies enabled the rapid accumulation of large TCR repertoire data. However, there are few efforts in integrating the growing single-cell immune profiling datasets across studies, and analyzing linked transcriptome and TCRα/β sequences spanning different individuals and diseases due to the technical batch effects across studies.

TCRs with high sequence similarity in CDR3 regions that were used to directly bind antigen peptides may indicate their relatively high chance in recognizing the same antigen peptide. Accordingly, various computational tools have been developed to perform unsupervised TCR clustering mostly based on CDR3 amino acids. Based on sequence similarity algorithms, GLIPH/GLIPH2, TCRdist/TCRdist3, iSMART, GIANA, and ClusTCR have been proposed.^20–26^ TCRdist3 and ClusTCR could take CDR3α and CDR3β amino acids as input, users could also provide CDR1 and CDR2 sequences as optional.^23,26^ While iSMART focuses on CDR3β amino acids, TRBV gene usage information could also be supplied as input.^24^ GIANA not only considered the sequencing similarity but also amino acid physicochemical properties.^25^ However, it can only take fixed length CDR3β amino acid sequences as input which greatly limited its general application.^25^ GLIPH2 could be applied to variable length CDR3α/β as well as TRBV gene usage simultaneously, and thus have been the most widely used tool in this field.^21^ The above-mentioned algorithms mostly based on clustering CDR3β amino acid information. Moreover, the computation time for most of these similarity-based algorithms increases non-linearly, which hinders their applications on large million-scale TCR data. Recently, deep-learning based methods have also been proposed especially for TCR-pMHC prediction tasks including DeepTCR, NetTCR-2.0, and TCR-BERT.^27–29^ Even though those methods were mainly developed for the prediction of TCR-pMHC binding, embeddings learnt from those methods could also be adapted to unsupervised TCR clustering with paired CDR3α/β sequences. ^27–29^ Aside from TCR sequence similarities, T cell states also contribute significantly to the identification of potential functional TCRs. Simultaneously capturing transcriptome and full-length paired TCR information from single-cell immune profiling dataset would thus substantially enhance our capability to identify functional TCR with right antigen specificity and the proper cell state. CONGA and Tessa both incorporate transcriptome information in their model to jointly represent TCR and cell state similarity.^30,31^ However, Tessa only combined CDR3β with transcriptome information, while CONGA used kernel PCA (Principal Component Analysis) for the transcriptome representation, limiting its scalability for large million-level dataset analysis and its extension for incorporating with other datasets due to potential batch effects.

In order to address major challenges mentioned above, we first collected million-level single-cell immune profiling datasets for T cells with various disease conditions including solid tumor, leukemia, inflammation/autoimmune, infections and healthy with full-length TCRα/β chains, single-cell transcriptome, HLA (Human Leukocyte Antigen) genotype and disease meta-information. With this massive dataset, we systematically analyzed the VJ gene preference and amino acid preferences in CDR regions for different CD4, CD8 cell subtypes. In addition, we discovered in general higher coherence for TCRα and TCRβ chains in antigen-experienced CD8 T cells. We also detected the existence of cross-individual public TCRs for both TCRα and/or TCRβ chains and characterized the general features for public TCRs. With this extensive reference atlas, we also developed a deep-learning based framework called TCR-DeepInsight to jointly represent similarity for TCR and transcriptome information. By incorporating the disease specificity scoring system proposed and the reference atlas established in this study, TCR-DeepInsight would be capable of identifying biologically meaningful disease-associated TCRs (dTCR) clusters effectively.

## RESULTS

### Integration of million-level paired TCRa/β repertoire with single-cell transcriptome information

In recent years, the advent of high-throughput single-cell sequencing technologies has enabled the profiling of immune repertoires at unprecedented depth and resolution. We previously developed a database called human Antigen Receptor database (huARdb).^32^ With our continuous efforts, these data have been expanded to 28 studies and 464 biological samples from 291 individuals, 9 tissue types, and 20 disease conditions including solid tumor, leukemia, inflammation/autoimmune, infections and healthy (**Figure1A and 1B; TableS1**). Altogether, we have collected 1,017,877 high-confidence T cells with paired transcriptome and full-length α/β TCRs information, which enable us to perform a large and unbiased analysis of the paired TCRα/β repertoire across diseases and individuals. These datasets contain 459,013 unique TCRα chains defined by the same TRAV(T cell receptor alpha variable gene)-CDR3α-TRAJ(T cell receptor alpha joining gene), and 604,728 unique TCRβ chains defined by the same TRBV(T cell receptor beta variable gene)-CDR3β- TRBJ(T cell receptor beta joining gene) (**Figure1A**). We defined clonally expanded T cells as T cells with the same V/J genes and CDR3 amino acid sequence (consist of TRAV/TRBV motif, CDR3α/β middle region (mr), TRAJ/TRBJ motif) of both α and β chains in the same individuals (**FigureS1A**). After merging clonal expanded TCRs, we obtained 635,393 unique TCR α/β clonotypes from 291 individuals. In addition, we also extracted the HLA genotypes for each individual from the single cell transcriptome dataset using arcasHLA.^33^ For all the 291 individuals collected in this study, A*02, B*15, C*07 were the most frequent MHC class I molecules, while DRB*103, DRB*115, DQB*103, DQA*101, DPB*102, DPB*104 were the most enriched MHC class II molecules (**FigureS1B-S1H**).

**Figure 1.**
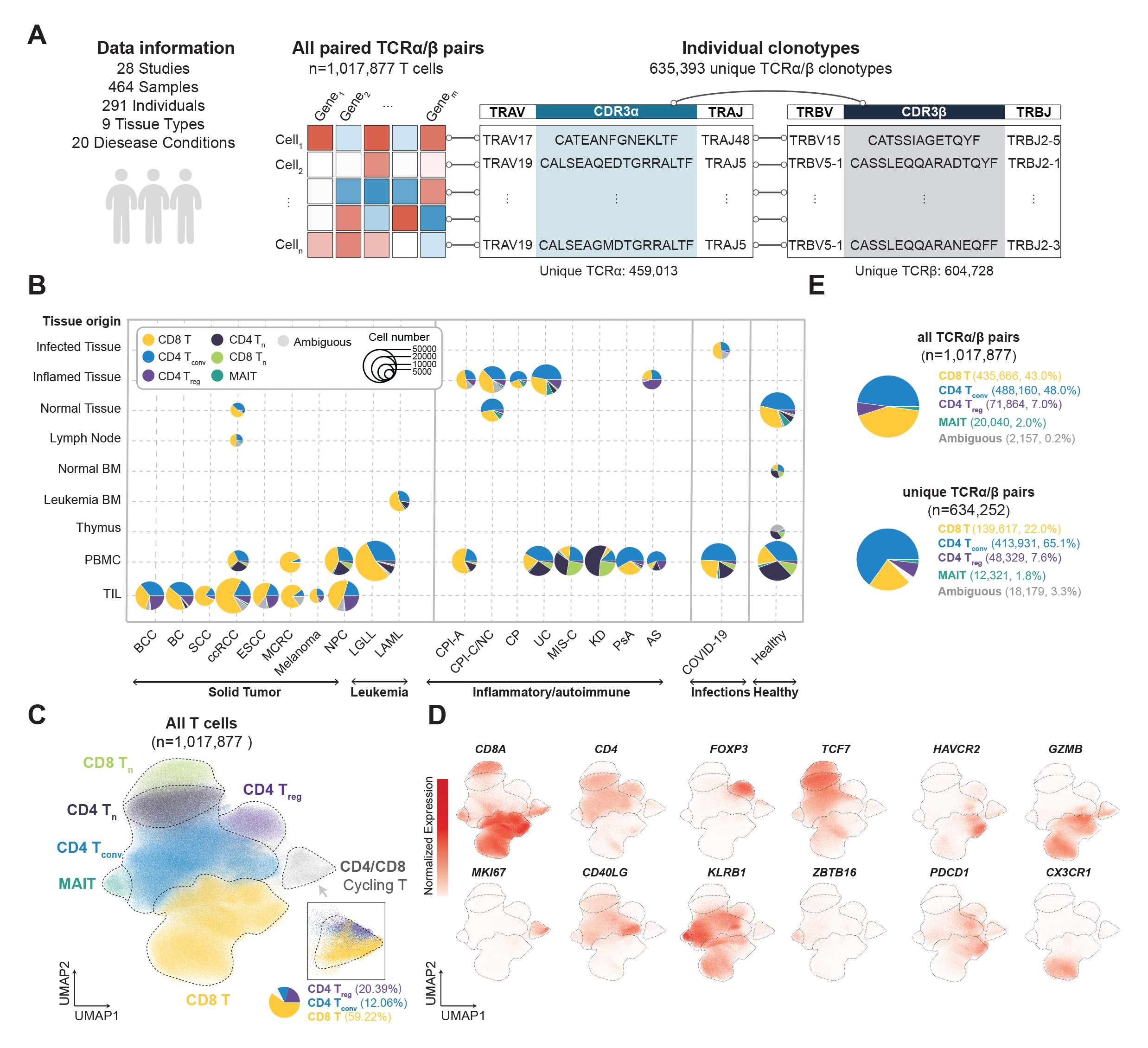
Overview of a million-level single-cell paired transcriptome and full-length TCR sequences dataset. **(A)** Schematic representation of the joint single-cell RNA sequencing (scRNA-seq) and full-length single-cell T cell receptor sequencing (scTCR-seq) data collected in this study. **(B)** Pie chart illustrating the distribution of T cell lineages from different tissue origins and disease types in the collected data. **(C)** Uniform manifold approximation and projection (UMAP) representation and cell lineage annotation for variational autoencoder (VAE)-based data integration for million-level scRNA-seq data. The panel in the bottom right corner shows a zoomed-in UMAP for cycling CD4 T_conv_, CD8 T, and CD4 T_reg_ cells. **(D)** Feature plot for key marker genes, including *CD8A, CD4, FOXP3, TCF7, HAVCR2, GZMB, MKI67, CD40LG, KLRB1, ZBTB16, PDCD1*, and *CX3CR1*, used for cell lineage annotation. **(E)** Pie chart depicting the composition of cell lineages for all TCRα/β pairs (upper panel) and unique TCRα/β pairs (lower panel). CD4 T_conv_: conventional CD4 T cells; CD4 T_reg_: regulatory CD4 T cells; CD4 T_n_: naïve CD4 T cells; CD8 T_n_: naïve CD8 T cells; MAIT: mucosal-associated invariant T cells. BCC: basal cell carcinoma; BC: breast cancer; SCC: squamous cell carcinoma; ccRCC: clear cell renal carcinoma; ESCC: esophagus squamous cell carcinoma; MCRC: metastatic colorectal cancer; BALL: B-cell acute lymphoblastic leukemia; NPC: nasopharyngeal carcinoma; LGLL: large granular lymphocytic leukemia; LAML: acute myeloid leukemia; UC: ulcerative colitis; CPI-C: checkpoint inhibitor associated colitis; CPI-NC: checkpoint inhibitor treatment with no colitis; CP: chronic pancreatitis; MIS-C: multisystem inflammatory syndrome; KD: Kawasaki disease; CPI-A: checkpoint inhibitor associated arthritis; AS: ankylosing spondylitis; PsA: psoriatic arthritis. BM: bone marrow; PBMC: peripheral blood mononuclear cell. TIL: tumor-infiltrating lymphocyte.

To gain an accurate cellular phenotype associated with full-length TCRs from gene expression profile, we first applied a rigorous approach to eliminate batch effects across samples and studies involving millions of single cells. We integrated the gene expression datasets with an in-house developed variational autoencoder (VAE) based deep-learning framework, and obtained a latent embedding of transcriptome features representing T cell subtype and functional states (**Figure1C**). We manually categorized T cells into six major states including, naïve (CD8 T_n_) and antigen-experienced CD8^+^ T cells (CD8 T), naïve (CD4 T_n_), conventional memory (CD4 T_conv_) and regulatory CD4^+^ T cells (CD4 T_reg_), and Mucosal invariant T cells (MAIT), aligned with the expression pattern of key marker genes (**Figure1C and 1D**). T cells including CD8 T, CD4 T_conv_ and CD4 T_reg_ in cycling status also formed a unique cluster in the integrated UMAP, suggesting the distinctive transcriptome features in cycling T cells (**Figure1C and 1D**). Using our VAE model, we successfully integrated data from 28 studies with minimal batch effects, and different cell subtypes were evenly distributed across various studies (**FigureS2A**). We observed a diverse cell type composition across solid tumor, inflammation, CPI-irAE (checkpoint inhibitor associated immune-related adverse events), AML (Acute Myeloid Leukemia), COVID-19, T-LGLL (T-cell large granular lymphocyte leukemia), and healthy conditions (**Figure1B and S2B**). To validate our cell type annotation, we applied the singleR automated cell type annotation tool, using PBMC T cell subtypes from healthy donors as cell type annotation reference, and found consistent cell subtype annotation (**FigureS2C**). ^34,35^

In this dataset, the percentage of unique TCRα/β pairs derived from CD4 T cells is higher than the percentage of CD4 T cells, indicating overall higher clonal expansion level of CD8 T cells than CD4 T cells (**Figure1E**). Overall, all TCRα/β pairs collected in this study consisted of 43.0% CD8 T cells, 48.0% CD4 T_conv_, 7.0% CD4 T_reg_, 2.0% MAIT cells, and 0.2% ambiguous (transcriptome can not be classified into either CD4 or CD8 T cells, see **Methods**), while our unique TCRα/β pairs across all individuals (n=634,252) consisted of 22.0% CD8 T, 65.1% CD4 T_conv_ cells, 7.6% CD4 T_reg_, 1.8% MAIT cells, and 3.3% ambiguous (identical TCRα/β pairs linked with cells with no dominant state, see **Methods**) (**Figure1E**). So far, to our knowledge, this is the most comprehensive atlas containing millions of TCRα/β pairs with transcriptome, full-length paired-TCR, HLA, and disease information.

### Association of intrinsic TCR features and major T cell subtypes

Previous studies of TCR repertoire using high-throughput sequencing have revealed preferences in VJ usage selection, and significant bias in VJ segment joining has been identified in single-chain repertoires.^36^ By using single-cell TCR sequencing followed by CD4 and CD8 FACS sorting, TCRα/β pairing have been proposed as a more precise way to infer repertoire functionality.^37^ With our reference atlas containing paired full-length TCR and transcriptome profiles, we can perform comprehensive analysis of the intrinsic features of TCR in different cell subtypes.

We calculated the odds ratio and presented significantly enriched TRAV-TRAJ and TRBV-TRBJ joining and TRAV-TRBV pairing in either CD8 T, CD4 T_conv_, CD4 T_reg_, or MAIT cells (**Figure2A; TableS2**). Our results confirmed the well-recognized phenomena that MAIT cells prefer to select TRAV1-2 paired with TRAJ12, TRAJ20 and TRAJ33 to form its invariant α chains.^38^ For β chain, MAIT cells preferentially select TRBV6-4, TRBV4-2, TRBV6-1, TRBV20-1, and TRBV6-2 as reported before.^39^ With the pairing information for TCRα/β chains, we were able to identify the significant preference for TRBJ2-1 and TRBJ2-3 in TRAV1-2^+^TRAJ33^+^TRBV6-4^+^ MAIT cells.

**Figure 2.**
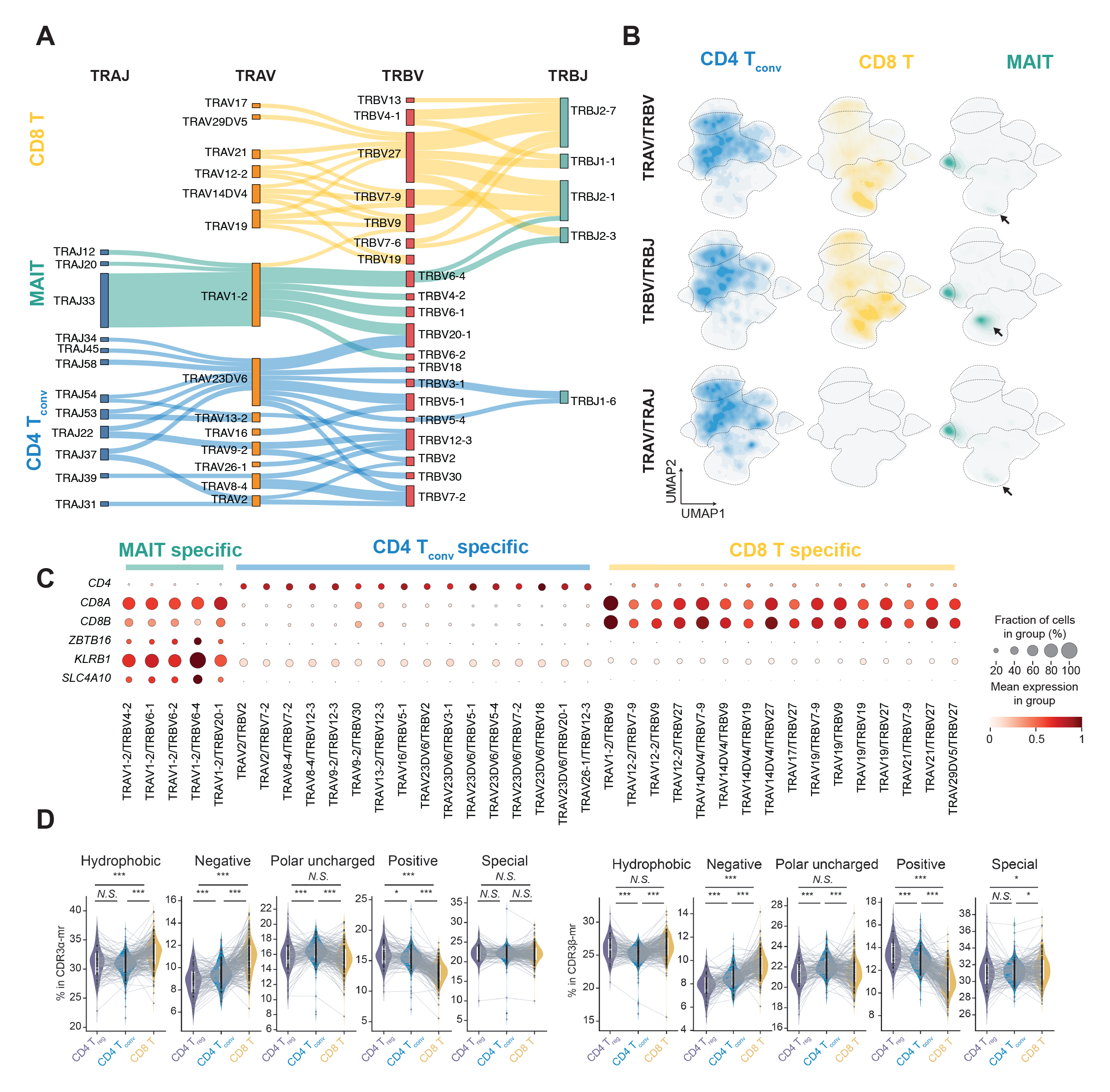
Analysis of T cell receptor (TCR) VJ segment pairing and amino acid usage in different T cell types. **(A)** Sankey plot illustrating the enriched TRAV/TRAJ, TRAV/TRBV, and TRBV/TRBJ pairing and joining preferences in CD8 T, MAIT, and CD4 T_conv_ cells. **(B)** Density plot showing the distribution of cells with enriched TRAV/TRAJ, TRAV/TRBV, and TRBV/TRBJ pairing in integrat-ed UMAP for CD8, MAIT, and CD4 T_conv_ cells. **(C)** Dot plot displaying the expression level of lineage-specific genes in CD8 T, MAIT, and CD4 T_conv_ cells with enriched TRAV/TRAJ, TRAV/TRBV, and TRBV/TRBJ. **(D)** Violin plot showing the preference for amino acids with different physicochemical properties in CDR3 middle region (mr) of CD8, CD4 T_conv_, and CD4 T_reg_ cells (*** *p-value* < 0.001, * *p-value* < 0.01, paired *t*-test; N.S.= not significant). CD8: conventional CD8 T cells; CD4 T_conv_: conventional CD4 T cells; CD4 T_reg_: regulatory CD4 T cells; MAIT: mucosal-associated invariant T cells; TRAV: TCR alpha variable; TRAJ: TCR alpha joining; TRBV: TCR beta variable; TRBJ: TCR beta joining. Each grey dot represent one individual, lines connecting grey dots indicate amino acid usage in different cell types within one individual. White boxes within violin plot represent the distribution from 25 percentile to 75 percentile.

Our analysis of TCR repertoire in CD8 T cells revealed no significant preference for TRAV-TRAJ genes. However, we identified TRBV27/TRBJ2-7, TRBV7-9/TRBJ2-1, and TRBV27/TRBJ2-1 as the top three significantly enriched TRBV/TRBJ joining pairs. In terms of TRAV/TRBV pairing, we found that TRAV19/TRBV7-9, TRAV14DV4/TRBV7-9, and TRAV14DV4/TRBV9 were the most dominant pairings (**Figure2A**).

The most enriched TRAV/TRBV pairings in CD4 T_conv_ cells were TRAV23DV6/TRBV20-1, TRAV23DV6/TRBV5-1, and TRAV8-4/TRBV7-2 (**Figure2A**). We observed significant enrichment in various TRAV/TRAJ joining pairs, with TRAV9-2/TRAJ22 being the most dominant pairing. In contrast, we detected only two significantly enriched TRBV/TRBJ joining pairs, namely TRBV3-1/TRBJ1-6 and TRBV5-4/TRBJ1-6 (**Figure2A**). Visualizing enriched TRAV/TRAJ, TRBV/TRBJ joining and TRAV/TRBV pairing enriched in CD8 T, CD4 T_conv_, or MAIT cells on the VAE integrated UMAP (Uniform Manifold Approximation and Projection) showed a diverse distribution with no specificity towards naïve or memory repertoire (**Figure2B**). Marker gene expression in cells with selected VJ gene usage further confirmed the TRAV/TRAJ, TRBV/TRBJ, and TRAV/TRBV preference in CD8 T, CD4 T_conv_, and MAIT cells (**Figure2C**). For example, MAIT-specific joining coincides with relatively high expression of MAIT-specific gene markers including *KLRB1* (Killer Cell Lectin-Like Receptor Subfamily B Member 1), *ZBTB16* (Zinc Finger and BTB Domain Containing 16), and *SLC4A10* (Solute Carrier Family 4 Member 10) (**Figure2C**). Interestingly, we also observed an enrichment of cells using MAIT specific VJ segments, which were not co-localized with conventional MAIT clusters on the integrated UMAP (**Figure2B**). These groups of cells were derived from a study from T-LGLL patients^40^, and displayed transcriptome features of terminal effector T cells including high expression of *CX3CR1* (X3-C Motif Chemokine Receptor 1) and *GZMB* (Granzyme B)(**Figure1D**). Although we do not know the origin of these T cells, nor whether they recognize MR1 (Major Histocompatibility Complex class I-related molecule 1)-restricted antigens as traditional MAIT cells or regular peptide antigens recognized by CD8 T cells, these findings are interesting and the functional role of these T cells in T-LGLL may require further investigation.

The CDR1 and CDR2 regions of the TCR typically contact the conserved α-helices of MHCs, and the usage of amino acids in these regions may influence the recognition between TCRs and class I and class II MHCs.^41^ Comparing to both CD4 T_conv_ cells and CD4 T_reg_ cells, our analysis indicated that CD8 T cells utilize significantly less hydrophobic and special amino acids, and more negatively, positively charged or polar uncharged amino acids in the CDR1α region (**FigureS3A**). For CDR1β region, no significant preferences for hydrophobic amino acids in CD4 T_reg_, CD4 T_conv_, or CD8 T cells, while significantly more negatively charged amino acids usage and less polar uncharged or special amino acids usage for CD8 T cells compared with CD4 T_reg_ and CD4 T_conv_ cells (**FigureS3B**). In CDR2α region, significantly more usage of polar uncharged and less usage of positively charged or special amino acids were detected for CD8 T cells comparing with CD4 T_reg_ or CD4 T_conv_ cells, while less hydrophobic amino acids and more negatively charged amino acids usage in CD4 T_reg_ comparing with CD8 T and CD4 T_conv_ cells (**FigureS3C**). Compared with both CD4 T_conv_ and CD4 T_reg_ cells, more hydrophobic and negatively charged and less polar uncharged, positively charged or special amino acids usage were observed in CD8 T cells for CDR2β regions (**FigureS3D**).

As the CDR3 region has been demonstrated as the most important part of the TCR in recognizing MHC class I or MHC class II displayed peptide, we asked whether the CDR3 repertoire may display different characteristics in various T cell subtypes, including CD4 T_reg_, CD4 T_conv_, and CD8 T cells. In our analysis, we find limited amino acid length difference for either CDR3α/β or CDR3α/β-mr in CD4 T_reg_, CD4 T_conv_, and CD8 T cells, while MAIT cells used shorter CDR3α or CDR3α-mr due to its relatively invariant α chain usage (**FigureS3E; TableS3**). Interestingly, we found that CD8 T cells use a significantly higher percentage of hydrophobic and negatively charged amino acids, and a lower percentage of polar uncharged and positively charged amino acids in the CDR3-mr for both α and β chain compared with CD4 T_reg_ and CD4 T_conv_ cells (**Figure2D and 2E; TableS4**). CD4 T_reg_ cells showed the lowest percentage of negatively charged amino acids and the highest percentage of positively charged amino acids in CDR3-mr for both chains compared with CD8 T and CD4 T_conv_ cells, while CD4 T_conv_ favored the usage of polar uncharged amino acids in CDR3-mr for both chains compared with CD8 T and CD4 T_reg_ cells (**Figure2D and 2E**).

### TCRa/β pairing is more coherent in memory T cells than in naïve T cells

As a comprehensive atlas of paired and full-length TCRα/β, our dataset is useful for understanding the V(D)J recombination biology and TCRα/β pairing patterns. As it has been previously reported that B cell receptors (BCRs) with the same heavy-chain-variable (VH) gene and CDRH3 (heavy-chain complementarity-determining region 3) amino acid sequence in their heavy chains from memory B cells or plasma cells that produce functional antibodies adopt the same light-chain-variable (VL) gene in their light chains.^42^ The phenomenon in memory B cells but not in naïve B cells is referred to as light chain coherence in BCRs.^42^ We thus hypothesized that a similar phenomenon may also exist in TCRs in memory state (CD4 T_conv_ cells or CD8 T cells). In our paired TCRα/β repertoire collections, we can detect TCRα/β chains using the same TRBV-CDR3β paired with the same TRAV gene but different CDR3α, or different TRAV gene and different CDR3α (**Figure3A**). In addition, we also observed TCRα/β chains with the same TRBV gene and similar but mismatched CDR3β pairing with either the same or different TRAV and different CDR3α (**Figure3A**). Similar cases were also observed when analyzing TRBV usage and CDR3β for relatively fixed TRAV-CDR3α (**Figure3B**).

**Figure 3.**
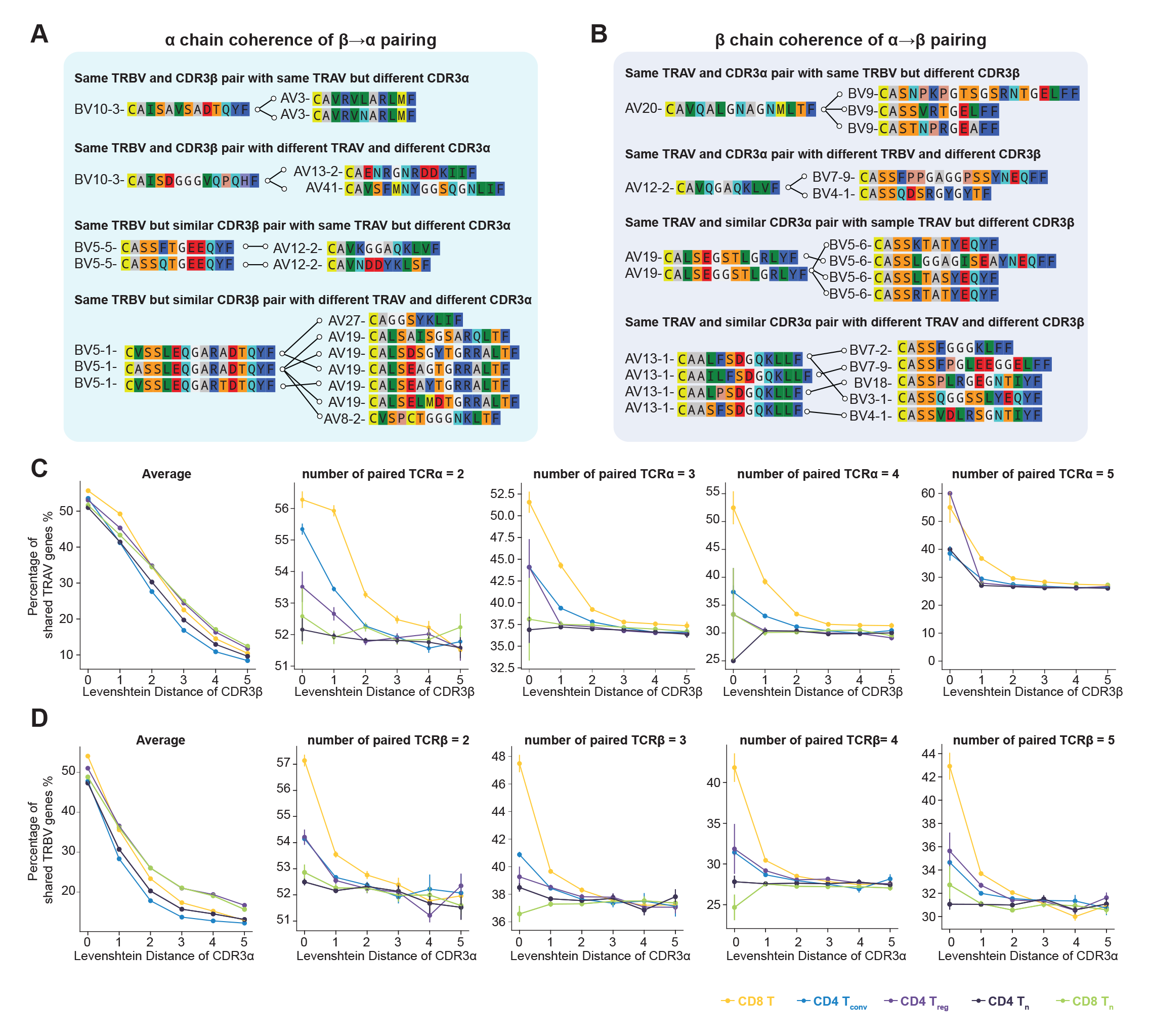
Coherence analysis of TCRα-TCRβ pairing. **(A, B)** Example of α coherence of β-α pairing (**A**) and β coherence of α-β pairing (**B**). T cells are grouped if (1) they had the same TRBV (α coherence) or same TRAV (β coherence); (2) they had the CDR3β (α coherence) or CDR3α (β coherence) amino acid sequence with Levenshtein distance differences less than 6; (3) they belong to the same T cell types including CD8 T, CD4 T_conv_, CD4 T_reg_, CD4 T_n_, and CD8 T_n_. **(C, D)** TRAV gene (**C**) and TRBV gene (**D**) usage for α coherence of β-α pairing (**C**) and β coherence of of α-β pairing (**D**) at different Levenshtein distance for CDR3α (**C**) or CDR3β (**D**) in different T cell types.

To analyze the TCRα/β chain coherence, we grouped TCRs with same TRBV by their deriving T cell subtypes (CD8 T_n_, CD8 T, CD4 T_n_, CD4 T_conv_, and CD4 T_reg_) and CDR3β similarity (CDR3β within 10 Levenshtein distance), and then calculated the percentage of TCRα/β pairs using the same TRAV gene. Our results suggested that TRAV coherence was in general higher in CD8 T cells in memory state than other cell types (**Figure3C**). Consistent phenomena were observed for grouping TCRs with the same TRAV and allowing mismatches in CDR3α, showing strongest TRBV coherence in CD8 T cells (**Figure3D**). Interestingly, when grouping TCRs by their CDR3β allowing more than two mismatches on average, we can no longer observe the TRAV coherence for CD8 T cells (**Figure3C**). However, TRBV coherence in CD8 T cells only allows one mismatch in CDR3α (**Figure3D**). The occurrence of the identical V gene usage in a randomly chosen set of TCRs would be a rare event. The increased V gene coherence observed in CD8 memory T cells might suggest these cells have experienced specific antigen. Identifying additional TCR clusters exhibiting V gene coherence could potentially lead to the discovery of TCRs that possess shared antigenic specificity.

### Characterization of public TCRs

Discovering TCRs with identical VJ segments and CDR3 amino acid sequences for both chains in different individuals is often considered an extremely rare event due to the total combinatorial diversity of TCR repertoire.^43^ These TCRs, hereafter called public TCRs, are thought to be generated by the selection of the same antigen epitope displayed by the same MHC molecule. In our massive paired TCRα/β collection, we identified 858 public TCRα/βs (with identical TRAV-CDR3α-TRAJ, and TRBV-CDR3β-TRBJ), 66,471 public TCRαs (identical TRAV-CDR3α-TRAJ), and 10,930 public TCRβs (identical TRBV-CDR3β-TRBJ) (**Figure4A; TableS5-S7**). Given the comparable number of total unique TCRα and TCRβ in the dataset, the significantly higher number of public TCRα indicates a generally more conserved α chain repertoire across populations. Cell states linking to public TCRα/β pairs were mostly enriched in CD8 T, CD4 T_conv_, and MAIT cells representing 35.8%, 36.0%, and 13.3% respectively, while in both public TCRα and TCRβ, CD4 T_conv_ were most dominant compared to other cell states (**Figure4B**). Public TCRs (either TCRα/β pairs, TCRα, or TCRβ) shared in only two individuals displaying variable cell states were classified into ambiguous (see **Methods**).

**Figure 4.**
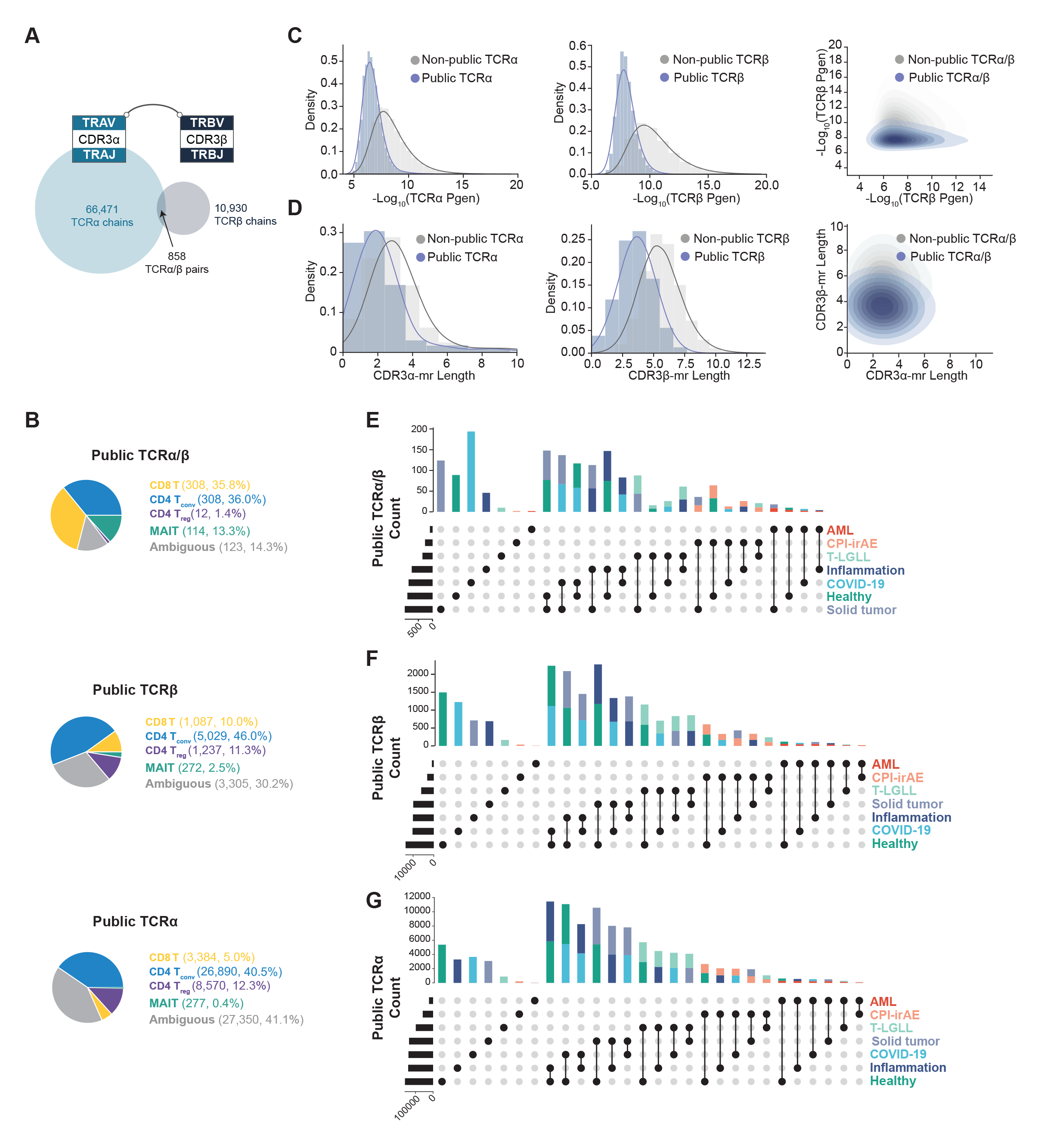
Public TCR analysis across samples and studies. **(A)** Venn diagram of public TCR defined by same α chain (same TRAV-CDR3α-TRAJ), β chain (same TRBV-CDR3β-TRBJ) or both chains. **(B)** Pie chart for cell type composition of public TCRα/β (upper panel), TCRα (middle panel), or TCRβ (lower panel). **(C)** Distribution of TCR generation probability for TCRα (left panel), TCRβ (middle panel), or TCRα/β (right panel) of public and non-public TCRs. **(D)** Distribution of CDR3 middle region length for TCRα (left panel), TCRβ (middle panel), or TCRα/β (right panel) of public and non-public TCRs. **(E-G)** Upset plot of disease composition of public TCRα/β (**E**), TCRα (**F**), or TCRβ (**G**).

To understand potential characteristics contributing to TCRα/β recurrence, we annotated each TCRα or TCRβ sequence with generation probability calculated by OLGA^44^ using CDR3 amino acid sequence and VJ segment genes. We found public TCRα and TCRβ, or public TCRα/β pairs were all associated with significantly higher generation probability (**Figure4C**). Compared to non-public TCRs, the negative log-transformed generation probability (-Log_10_P_gen_) is smaller in public TCRα (6.7067±0.0019 compared to 8.4534±0.0027) and TCRβ (7.9750±0.0050 compared to 10.3934±0.0025) (**Figure4C**). Interestingly, when visualizing generation probability for TCRα and TCRβ in public TCRα/β pairs, TCRβ instead of TCRα exhibited greater difference compared to non-public TCRα/β pairs (**Figure4C**).

We also analyzed CDR3-mr amino acid length in public TCRs, and observed a generally shortened CDR3mr amino acid length in either public TCRα (2.1834±0.0034 compared to 3.2468±0.0028) or public TCRβ (3.6881±0.0080compared to 5.5380±0.0021) (**Figure4D**). Consistent with the observation for generation probability, when examining the CDR3-mr amino acid length for public TCRα/β pairs, only TCRβ but not TCRα CDR3-mr amino acid length demonstrated a shortening pattern (**Figure4D**). This phenomenon implies that TCRβ may have a more significant role in TCR recurrence resulting from same antigen challenge, which aligns with previous understanding that TCRβ contributes more to antigen specificity determination.

### Identification of epitope-and disease-related public TCRs

We hypothesized that public TCRs that occur in different individuals in the same disease setting may bind to disease-specific epitopes. We asked whether public TCRα, TCRβ or TCRα/β pairs would be restricted in common diseases such as solid tumours, autoimmune diseases, or COVID-19. Most of the public TCRs are shared in individuals with different disease settings, and these TCRs may not target disease-specific epitopes (**Figure4E-4G**). However, we indeed identified an enrichment of SARS-CoV-2-specific public TCRα/β pairs (but not in public TCRα or TCRβ), and this phenomenon may be explained by the relatively restrained epitope peptide sequence derived from SARS-CoV-2 (**Figure4E**).

We next asked whether the public TCRs would bind epitopes derived from common viruses such as Epstein-Barr virus (EBV), Cytomegalovirus (CMV), and Influenza A virus that remain latently present in target cells after the primary infection. In order to match TCRs with known epitope specificity, we build a reference dataset containing paired CDR3α/β sequences with experimentally validated epitopes from various sources (**FigureS4A; TableS8**).^22,27,45–47^ Since CDR3α/β-epitope pairs datasets mostly collected from EBV, CMV, Influenza A, or SARS-CoV-2 viruses, matched TCRα/β pairs with identical CDR3α and/or CDR3β sequences were mostly enriched for antigen listed above and do not harbor significant disease specificity (**FigureS4B**).

We performed in-depth analysis on public TCRs that were clonally expanded in at least two individuals across studies. Linking transcriptome features to public TCRα/β defines their potential functional roles in different disease conditions. We identified public and clonally expanded TCRα/β pairs (TRAV5-CAESTGKLIF-TRAJ37/TRBV29-1-CSVGTGGTNEKLFF-TRBJ1-4 and TRAV21- CAVLMDSNYQLIW-TRAJ33/TRBV10-2-CASSEDGMNTEAFF-TRBJ1-1) in 12 or 7 HLA-A02:01 positive individuals, with matched epitope specificity to HLA-A02:01 restricted BMLF1280-EBV and LMP2-EBV by both CDR3α/β sequences, respectively (**FigureS5A**). We also find another public and clonally expanded TCRα/β pair (TRAV23DV6-CAASIGNFGNEKLTF-TRAJ48/TRBV4-2-CASSPSRNTEAFF-TRBJ1-1) in 5 HLA-B07:01 positive individuals matched to HLA-B07:02 restricted pp65-CMV epitope (**FigureS5A**). Public TCRα/β with previously unknown antigen specificity were also detected in our analysis. For example, TRAV26-2-CILRGAGGTSYGKLTF-TRAJ52/TRBV27-CASSLQGANYEQYF-TRBJ2-7 were identified in five HLA-B07* positive individuals, and TRAV17- CATEGDSGYSTLTF-TRAJ11/TRBV6-5 CASSGQGGYGYTF-TRBJ1-2 were detected in four HLA-A02* individuals (**FigureS5A**). T cells harboring these public TCRα/β pairs were likely to be clonally expanded and enriched in antigen-experienced CD8 T cells (**FigureS5A**).

Although observing public TCRs from individuals that are both clonally expanded and associated with the same disease are extremely rare events, we observed several such TCR pairs derived from CD4 or CD8 T cells. For example, the public and clonally expanded TCRα/β pairs enriched on antigen-experienced CD8 T cells (TRAV17-CATDYNQGGKLIF-TRAJ23/TRBV4-1-CASSQDPGAWETQYF-TRBJ2-5) were observed in two individuals with inflammatory diseases (ulcerative colitis and chronic pancreatitis) sharing HLA-A02, HLA-A03, HLA-B07, and HLA-C07 (**FigureS5B**). Public and clonally expanded TCR specific to psoriatic arthritis (TRAV17-CATDGYTGNQFYF-TRAJ49/TRBV12-3-CASSLGQNNEQFFTRBJ2-1) was also enriched with antigen-experienced CD8 T cells in two individuals sharing HLA-B15, and HLA-C03 (**FigureS5B**). We have also found other clonally expanded and public TCRs in either CD4 or CD8 T cells in solid tumors, acute myeloid leukemia, and COVID-19 associated with common HLAs (**FigureS5B**). These TCRs are highly likely to play crucial roles in various disease settings, although experimental validation remains necessary to confirm their importance.

### TCR-GEX joint-embedding learned by deep learning models facilitates TCRa/β clustering with disease-association and shared transcriptome state

The coherence in TCRα or TCRβ chains in antigen-experienced CD8 T cells is dependent on the CDR3α or CDR3β sequence with one or two mismatched amino acids. These results suggest that both CDR3α and CDR3β contributed to the forming of epitope specificity. In most cases, researchers were unable to link a TCR repertoire to a previously known set of antigen peptides. The MHC displayed epitope peptide for patients with cancer and autoimmune diseases, or even specific viral infections could be diverse across individuals, and so does the TCR repertoire. One of the tasks in TCR repertoire analysis is to cluster TCRs with higher probability to recognize the same antigen peptide. It has previously been discussed that using both CDR3α or CDR3β amino acid sequences could improve prediction accuracy in epitope specificity of TCRs.^48^ Moreover, it has been recently emphasized that incorporating TCR sequence and cell-specific covariates from single-cell data can enhance the inference of T cell antigen specificity.^49,50^

To robustly identify potential disease associated TCRα/β pairs considering both TCR sequence similarity and transcriptome features from million-level paired TCRα/β repertoire, we developed a deep-learning based framework named TCR-DeepInsight. We first extracted the latent embedding learned by the VAE model to represent the GEX features (**Figure5A**). In parallel, we adopted the Bidirectional Encoder Representations from Transformers (BERT) model to learn the underlying relationship between V genes and CDR3 sequence and the pairing pattern of the two chains (**Figure5B**). We used 635,322 unique TCRα/β CDR3 sequences along with TRAV, TRAJ, TRBV, TRBJ gene usage for pretraining BERT by randomly hiding certain VJ gene or amino acids used in either TCRα or β chains (**Figure5B**). The model does not require prior knowledge of the epitope binding specificity and thus clustering on the embedding of the output layer was unsupervised. Given the higher importance of TCRβ chain in its function, we apply the pre-trained model on the same dataset containing 635,322 unique TCRα/β pairs using the TCR embedding pooled from token features from TRAV, TRBV, CDR3β and TRBJ. After principal component analysis transformation to a 64 dimensional TCR embedding space, we found that the TCRs can be primarily clustered by TRBV usage in the 2-dimensional UMAP projection (**Figure5C**). This TCR embedding could be applied to downstream tasks including unsupervised TCRα/β clustering and query for TCRα/β with known epitope specificity (**Figure5C**). To match GEX embedding with TCR embedding and avoid single TCR embedding shared by multiple GEX embedding, we averaged the VAE latent embedding for unique TCRα/β pair by their deriving cells and obtained an aggregated GEX embedding (**Figure5C**). TCRα/β linked with multiple cell states were classified as the most dominant cell state, while TCRα/β linked cells with no dominant state were classified as ambiguous (**Figure5C**). Consistent with the original GEX embedding, similar cell states were clustered together on the UMAP of aggregated GEX embedding (**Figure5C**). We then concatenated the TCR embedding learned from BERT and the aggregated GEX embedding from VAE to obtain a TCR-GEX joint representation of TCRα/β pairs from million-level paired TCRα/β repertoire with single-cell transcriptome information (**Figure5C**). We found that our TCR-GEX joint representation was capable of capturing TCR similarity of amino acids for both CDR3α and CDR3β and a positive correlation between Levenshtein distance of amino acid sequence and Euclidean distance of the joint representation within eight Levenshtein distance (**Figure5D and 5E**).

**Figure 5.**
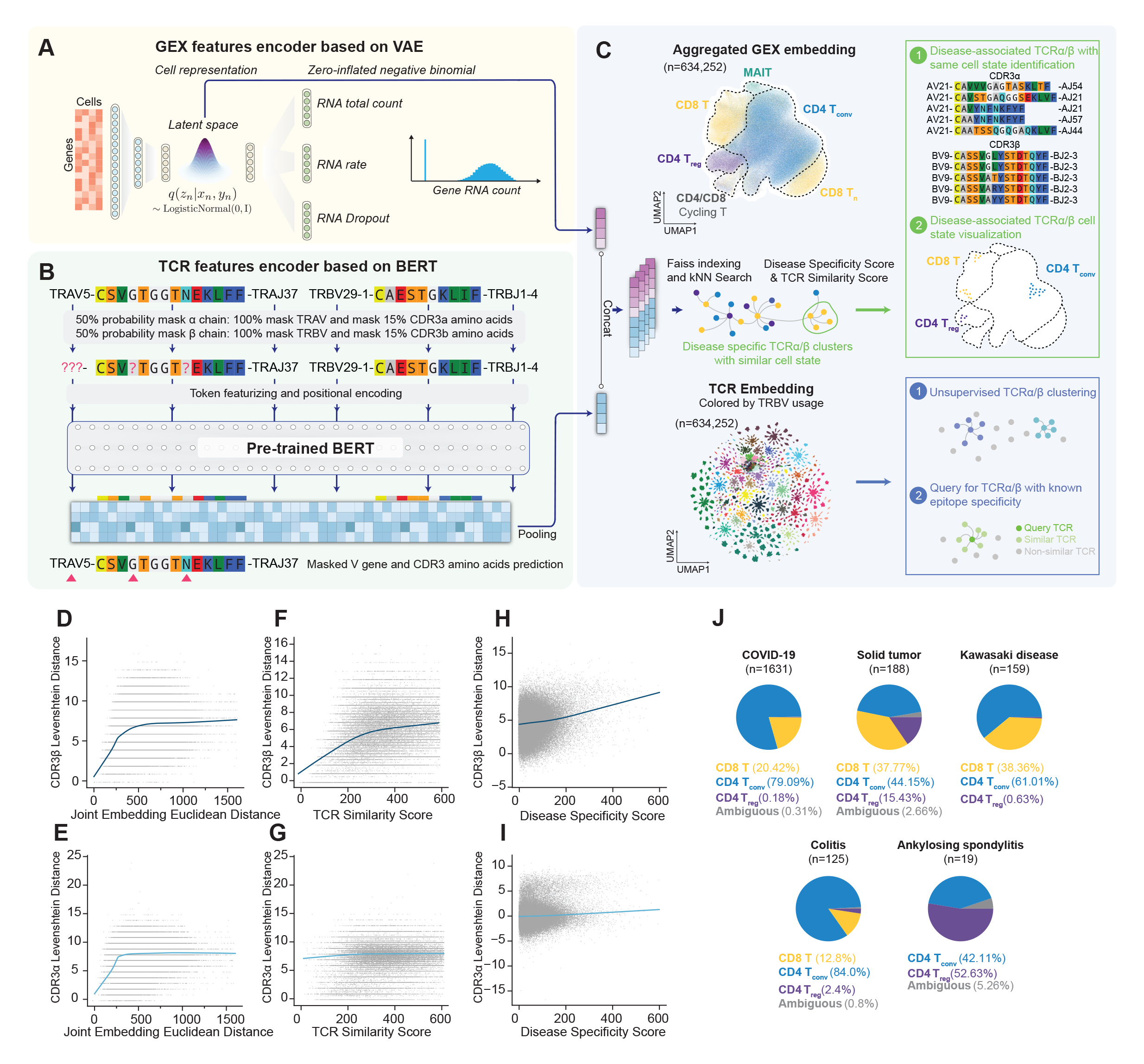
Integrating TCR sequences and transcriptome features to identify dTCR clusters with transcriptome similarity. **(A-C)** Schematic illustration for TCR-Deeplnsight model including VAE model used in transcriptome features extraction (**A**), TCR pre-trained with BERT model used in TCR features extraction (**B**), TCR embedding used for down-stream tasks including unsupervised TCR clustering, query TCR with known epitope specificity (**C**), and concatenated joint embedding with both TCR and transcriptome features (TCR-GEX joint embedding) used for downstream dTCR clusters identification, and GEX visualization (**C**). **(D, E)** Relationship of Levenshtein distance of amino acids in CDR3α (**D**) or CDR3β (**E**) and Euclidean distance of TCR-GEX joint embedding between random selected pairs of TCRs. **(F, G)** Relationship of average pair-wise Levenshtein distance of amino acids in CDR3α (**F**) or CDR3β (**G**) and the TCR similarity score of TCR-GEX joint embedding for all potential disease TCR clusters. **(H, I)** Relationship of difference in Levenshtein distance of amino acids in CDR3α (**H**) or CDR3β (**I**) and disease-specificity score of TCR-GEX joint embedding for all potential disease TCR clusters. **(J)** Pie-chart represent the cell type composition of dTCR clusters in different disease conditions.

With the massive paired TCRα/β repertoire along with disease information collected in this study, we were able to identify disease associated TCRα/β clusters with shared transcriptome state with TCR-DeepInsight (**Figure5C**). Using the Euclidean distance for unique TCRα/β pairs in TCR-GEX joint representation (referred as TCR distance), we clustered each TCRα/β pairs with 39 most similar TCR neighbors using GPU(Graphics Processing Unit)-accelerated indexed kNN (k-Nearest Neighbors, k=40) search (**FigureS6A**). Then we iteratively selected a unique TCRα/β pair as anchor and ranked its neighboring TCRs by its TCR-GEX joint representation similarity (**FigureS6A**). If an anchor TCR and its top similar neighboring TCRs (minimal 2) were derived from the same disease, these TCRs were then defined as disease-TCR-cluster, while the same count of next ranked TCRs were defined as non-disease-TCR-cluster (**FigureS6A**). TCR distance within disease clusters were then calculated and defined as TCR similarity score, while TCR distance between disease-TCR-cluster and non-disease-TCR-cluster were defined as disease specificity score (**FigureS6A**). Interestingly, we found a positive correlation between the TCR similarity score and the averaged Levenshtein distance of CDR3β but not CDR3α, while a positive correlation was found in the averaged between-cluster difference Levenshtein distance of CDR3β and the disease specificity score (**Figure5F-5I**). For each disease condition, we linked each TCR cluster with cell state information and calculated the TCR similarity score and disease specificity. Consistent with previous definition, disease-associated TCR clusters with dominant cell state were classified into either CD8 T cells, CD4 T_conv_, CD4 T_reg_, MAIT, or ambiguous. TCR clusters in each disease condition were then classified as dTCR clusters under certain cut-off (**FigureS6A**, see **Methods**). With the GEX information available, we were able to project dTCR clusters onto aggregated GEX and original GEX embedding to visualize their transcriptome features and additional information such as HLA information and clonal expansion level could also be retrieved (**FigureS6A**). Overall, we identified dTCR clusters in various disease conditions, including solid tumors, COVID-19, and Colitis containing relatively large sample numbers, while dTCR clusters could also be detected in Kawasaki disease and Ankylosing Spondylitis which have relatively fewer individuals (**Figure5J; TableS9**).

### Identification of dTCR clusters from various disease conditions

In solid tumors, one of the top dTCR clusters ranked by the TCR similarity score contains was derived from three A*33:85 and B*58 positive individuals with nasopharyngeal carcinoma (NPC), breast cancer (BC), and esophagus squamous cell carcinoma (ESCC). This dTCR cluster showed utilization of TCRα/β using TRAV17, TRAJ45, TRBV18 and TRBJ2-1, with a remarkably similar pattern of CDR3α and CDR3β amino acid sequences and no known antigen specificity have been reported (**Figure6A**). Another dTCR cluster associated with basal cell carcinoma (BCC) and breast cancer (BC) utilize TRAV29DV5, TRBV25-1 and TRBJ2-5 with similar CDR3 patterns. Both solid tumor dTCR clusters displayed coherence in the transcriptome states in antigen-experienced CD8 T cells as visualized by the projection to the aggregated and original GEX embedding (**Figure6A**). Interestingly, we also detected a CD4 T_conv_ dTCR cluster with extremely similar CDR3α/β sequences associated with NPC, BCC and SCC (squamous cell carcinoma), shared in 4 individuals with DQA*102 and DRB*107 (**FigureS6B**).

**Figure 6.**
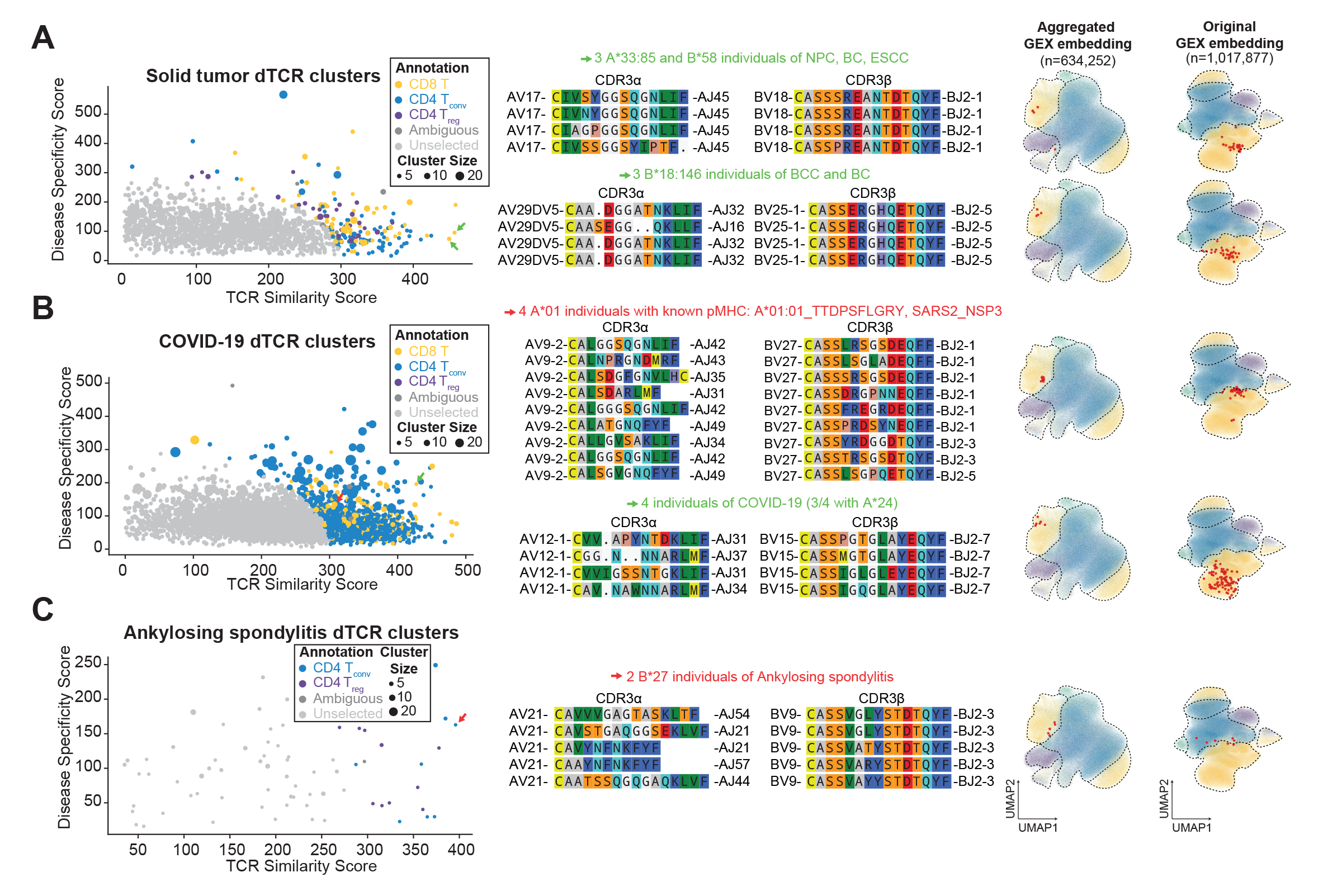
Representative dTCR clusters with transcriptome similarity in solid tumor, COVID-19 and ankylosing spondylitis. **(A-C)** Representative dTCR clusters of solid tumor **(A)**, COVID-19 **(B)**, and Ankylosing spondylitis **(C)**. Scatter plot of TCR similarity score, disease specificity score, and cell type annotation for dTCR clusters were shown (left panel), VJ gene usage and CDRα/β amino acid sequences were for selected dTCR clusters were shown (middle panel), and their transcriptome featurs on original and aggregated GEX embedding were visualized (right panel). Green arrows on scatter plot indicate dTCR clusters selected for visualization with no known antigen specificity, red arrows on scatter plot indicate dTCR clusters selected for visualization with previously reported antigen specificity. Red stars on aggregated and original GEX UMAP indiate the cells harboring these dTCRs.

We also identified dTCRs derived from COVID-19 patients with high disease specificity and transcriptome similarities. We found a dTCR cluster from 4 A*01 individuals utilizing TRAV9-2, and TRBV27 with similar CDR3β but not CDR3α sequence, whose deriving cells were clustered in the antigen-experienced CD8 T state (**Figure6B**). We found that this TCR cluster may recognize the A*01:01 restricted QYIKWPWYI derived from the nonstructural protein of SARs-CoV-2 reported by Francis et al.^47^ using tetramer staining followed by single-cell sequencing. We also found other COVID-19 dTCR clusters with high disease specificity, overlapping HLA alleles, and shared transcriptome states, but without previously known binding SAR-CoV-2 derived epitopes (**Figure6B**).

Our method is also applicable to datasets with few individuals. In a study containing two B*27 positive individuals with Ankylosing Spondylitis, we found that the top-ranked TCR cluster utilized TRAV21, TRBV9, and TRBJ2-3 with conserved CDR3β motif but diverse TRAJ usage and CDR3α sequence (**Figure6B**). It has been previously reported as an over-representation of TRBV9-CDR3-TRBJ2-3 sequence motifs in individuals with ankylosing spondylitis (AS) compared to healthy individuals who carry the B*27 allele and further research demonstrates that these TCRs recognize both self-antigen and microbial antigen displayed by the B*27.^51^ Cell types associated with these TCRs were located on the border of CD8 T cells, and CD4 T_conv_ cells in both aggregated and original GEX embedding, suggesting its potential functional role (**Figure6C**).

### Extension of TCR-DeepInsight for TCR query with known antigen specificity and model transfer on new dataset

TCR-DeepInsight provided a pre-trained BERT model on massive TCRα/β pairs and the dataset collected in this study provided a comprehensive reference TCR repertoire along with single cell transcriptome information. In order to infer potential antigen specificity for TCRα/β pairs collected, we next extended the TCR BERT pre-trained model to the task of querying TCRα/β with known epitope information. By processing TCRα/β pairs containing VJ gene usage and CDR3α/β sequences with known epitope information through our BERT pre-trained model, we obtained the query TCR embedding with known epitope information. Query TCR embedding and reference TCR embedding were then aligned and kNN search for epitope specific TCRα/β were performed. Under certain cut-off for TCR distance, TCRα/β in the reference dataset similar to query TCRα/β with known epitope information could be identified, and additional information such as HLA information, clonal expansion level and visualization on GEX embedding could be retrieved (**Figure7A**, see **Methods**). By querying two TCRα/β pairs recognizing A*24:01 restricted epitope NYNYLYRLF from SARS-CoV-2 Spike protein and an B*15:01 restricted epitope RVAGDSGFAAY from SARS-CoV-2 membrane glycoprotein, we identified TCR using similar VJ genes and CDR3β. For both queried TCRs, the top three most similar TCRs were both derived from COVID-19 disease condition, and cells harboring these TCRs predominantly were antigen-experienced CD8 T cells (**Figure7B, C**).

**Figure 7.**
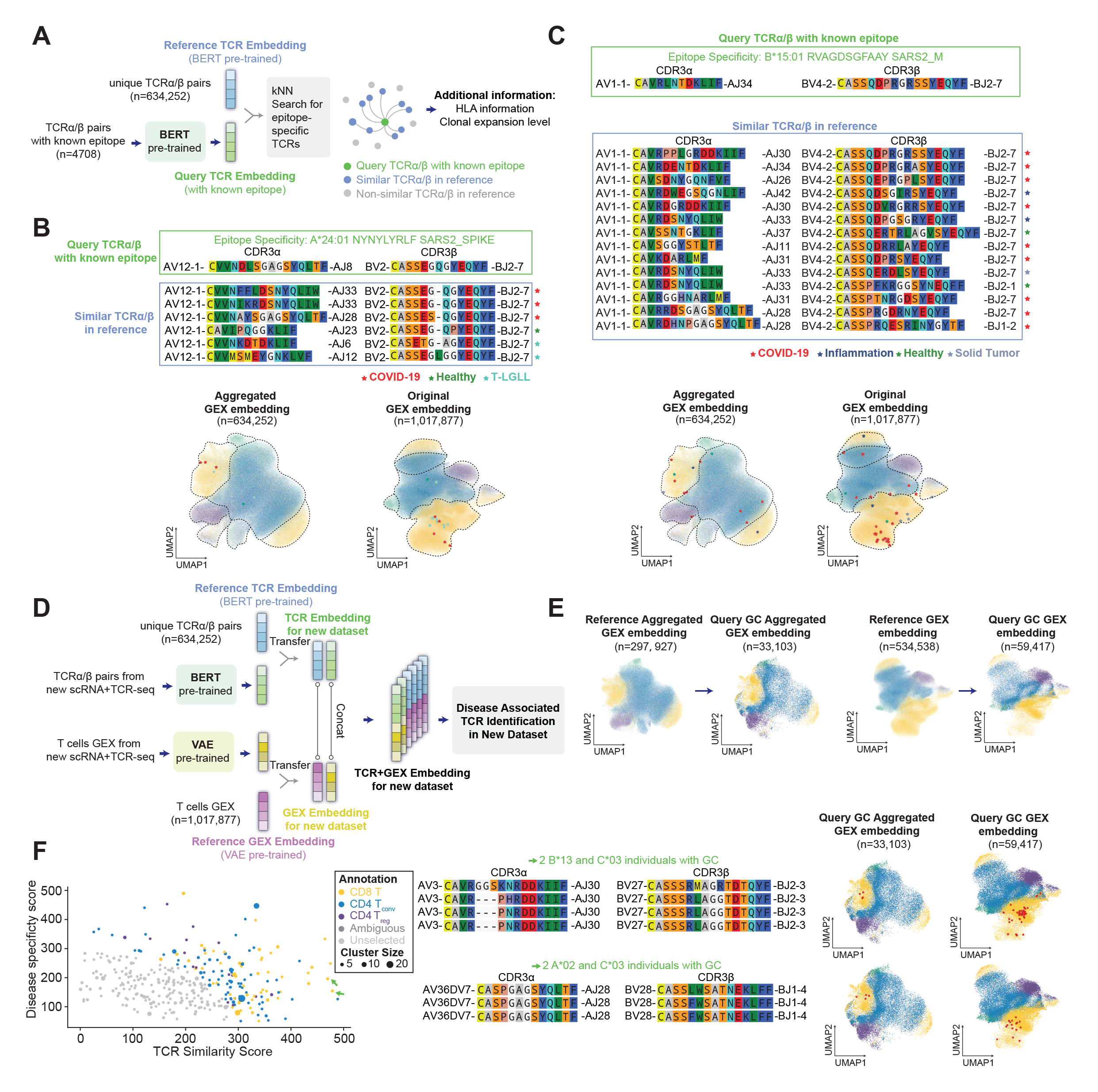
TCR-DeepInsight extended for query TCR with known epitope specificity and transfer on new dataset. **(A)** Schematic illustration for TCR-DeepInsight module used for querying TCRs with known epitope specificity in the reference TCR atlas. **(B, C)** Representative TCR cluster from the reference TCR atlas queried by an A*24:01 restricted epitope NYNYLYRLF from SARS-CoV-2 Spike protein (**B**) and an B*15:01 restricted epitope RVAGDSGFAAY from SARS-CoV-2 membrane glycopro-tein (**C**). Colored stars on aggregated and original GEX UMAP indiate the cells harboring these TCRs similar to the queried TCRs. Color correspond to disease conditions associated with TCRs. **(D)** Schematic illustration for transfering BERT pre-trained TCR model and VAE pre-trained GEX model to new scRNA-seq and scTCR-seq dataset to identify dTCRs. **(E)** UMAP representation of original GEX and aggregated GEX embedding for queried scRNA-seq and scTCR-seq dataset from gastric cancer and randomly down-sampled reference atlas. **(F)** Representative dTCR clusters identified in gastric cancer. Scatter plot of TCR similarity score, disease specificity score, and cell type annotation for dTCR clusters were shown (left panel), VJ gene usage and CDRα/β amino acid sequences were for selected dTCR clusters were shown (middle panel), and their transcriptome featurs on original and aggregated GEX embedding were visualized (right panel). Green arrows on scatter plot indicate dTCR clusters selected for visualization with no known antigen specificity. Red stars on aggregated and original UMAP indiates the cells harboring these dTCRs.

In general, TCR-DeepInsight provided a comprehensive reference containing millions of TCRα/β pairs with side-by-side transcriptome and disease meta-information, allowing more precise identification of disease-associated TCRs for novel datasets. To demonstrate the model transfer performance for TCR-DeepInsight on new dataset, we leveraged an additional scRNA-seq and scTCR-seq dataset to identify unique clusters of TCRα/β pairs associated with gastric cancer.^52^ Our method involved transferring the GEX embedding from the VAE model and the TCR embedding from the BERT model to new datasets. We accomplished this by implementing architecture surgery on the VAE model, a technique that allows the transfer of previously trained weights on a larger reference atlas to new smaller datasets (**Figure7D**). Simultaneously, the query TCR embedding can be obtained from the same pretrained BERT model as used in the reference dataset followed by PCA transformation using the same weight (**Figure7D**).

We observed that our query gastric cancer dataset could be projected into a common low dimensional space with our reference atlas using both unaggregated and aggregated GEX embeddings (**Figure7E**). By applying the same strategy as for other disease conditions, we identified two dTCR clusters associated with gastric cancer that exhibited high sequence similarity and disease specificity (**Figure7F; TableS10**). Notably, one TCR cluster derived from CD8 T cells and was clonally expanded, characterized by the usage of TRAV3, TRAJ30, TRBV27, and TRBJ2-3 genes, and CDR3α and CDR3β sequences that were similar in two individuals who were positive for B*13 and C*03 (**Figure7F**). The other TCR cluster derived from a more similar transcriptome state of CD8 T cells and was also clonally expanded, characterized by the usage of TRAV36DV7, TRAJ28, TRBV28, and TRBJ1-4 genes, and CDR3α and CDR3β sequences that were similar in two individuals who were positive for A*02 and C*03 (**Figure7F**).

## DISCUSSION

With the advent of single-cell immune profiling technology, vast amounts of single-cell gene expression (GEX) and T cell receptor (TCR) data have been generated. These data offer tremendous potential for studying TCR biology and identifying potential functional TCRs. However, a major challenge is the lack of a reference atlas that provides easy access to these datasets. On the other side, with the rapid development of TCR engineering, the use of TCR-T in disease immunotherapy is becoming increasingly promising. Single-cell immune profiling datasets provide a valuable platform for the identification of functional TCRs. However, the lack of efficient computational tools to integrate and identify functional TCRs remains a significant bottleneck in this field.

In this study, we established a comprehensive dataset comprising millions of T cells obtained from single-cell immune profiling, along with their single-cell transcriptomes, full-length paired TCRs, HLA types, and disease information. This dataset serves as a valuable reference atlas for a wide range of computational tasks in immunology. Using this massive dataset, we comprehensively analyzed the intrinsic features of TCRs in CD4 and CD8 lineages, identified coherence of public TCRα/β chains in memory CD8 T cells, and illustrated the widely-existing public TCRs across individuals. Moreover, we developed TCR-DeepInsight, a powerful and scalable deep-learning framework that can effectively cluster TCRα/β pairs with high transcriptome similarity in single-cell immune profiling datasets. With the disease scoring system embedded in TCR-DeepInsight, researchers can readily identify disease-associated TCR clusters using the dataset collected in this study as a reference. This tool offers a valuable resource for the analysis of single-cell gene expression and TCR data, facilitating the identification of disease-specific TCR repertoires in various disease contexts.

Our results highlight the potential of integrative analysis with large cross-samples datasets to gain deeper insights into T cell biology. Additionally, our reference atlas and TCR-DeepInsight tool offer versatile resources for immunologists to identify potential functional TCRs from single-cell immune profiling datasets. Most importantly, the dTCR clusters identified in this study, along with future potential dTCRs characterized with single-cell immune profiling and TCR-DeepInsight, may guide the design of personalized TCR-T (T cell receptor-engineered T cells) for precise treatment of various diseases, including cancers and autoimmune disorders, in the future.

We identified significant biases in the selection of VJ segment and amino acid usage in the CDRs of the TCRs associated with different T cell types. A restricted VJ segment usage was found for both TCR α and β chains in MAIT cells by virtue of their evolutionarily conserved recognition of MHC-like molecule MR1.^53^ We observed significant enrichment for TRAV/TRAJ, TRBV/TRBJ joining, and TRAV/TRBV pairing in CD8 T, CD4 T_conv_, and MAIT cells, which is in line with previous findings.^36, 37^ Additionally, a significant bias was observed in the usage of amino acids in the CDR1 and CDR2 loops. One possible explanation of these observations is that the TRAV/TRBV usage and the composition of amino acid usage after V(D)J recombination predispose TCRs to interact with MHC I or MHC II molecules.^41^ While no obvious bias for VJ segments were detected in CD4 T_reg_, we detected significant differences for the amino acids usage in CD4 T_reg_ including the preference for hydrophobic amino acids in CDR3β compared with CD4 T_conv_ as reported before.^54, 55^ These findings revealed the intrinsic molecular features difference between MHC class I and II recognition by TCR, and provide evidence supporting the plausibility of classifying CD4 or CD8 TCRs based on VJ gene usage and TCR amino acid sequence solely.

Given the immense diversity of TCRs, observing public TCRs with identical α and β chains is considered an extremely rare event. These TCRs are believed to share antigen specificity. Previous analysis on public TCRs have shown that TCR sequences are shared across multiple individuals, indicating a potential shared antigenic exposure.^56,57^ We comprehensively analyzed public TCRs for both α and β chains, or single chain, and found that public TCR sequences are generally shorter in the middle region of CDR3 for both α and β chains and have a higher generation probability, which is consistent with TCRs in mice.^58^ We also observed an enrichment of SARS-CoV-2-specific public TCRα/β pairs and identified public and clonally expanded TCRs with known antigen specificity for viruses such as EBV, CMV, and Influenza A. Meanwhile, we also identified numerous public TCRs shared in solid tumors and inflammation without known epitope specificity, which may imply their roles in recognizing tumor neoantigens or self-antigens. Coherent TCRα/β pairs are also considered to recognize similar antigens. By analyzing a comprehensive atlas of paired and full-length TCRα/β from different T cell subtypes, we found that TRAV coherence, or the tendency for TCRα chains to use the same TRAV gene when paired with the same TRBV-CDR3β, was generally higher in memory CD8 T cells than in other cell types. However, both the public TCR and coherence analysis focused on VJ gene usage and TCR sequence similarity without considering the potential cell transcriptome status, which may not be robust. Hence, developing new methods that can integrate both TCR and GEX data may help facilitate the identification of functional TCRs.

VAE has previously demonstrated its general applicability in scRNA-seq data feature extractions and batch effect removal across samples.^59^ Meanwhile, the natural language processing large-language model BERT has also been adapted in representation learning for DNA, protein, and TCR sequences.^29,60,61^ Deep-learning based framework allows representation learning for both transcriptome and TCR information. VAE based deep-learning framework in TCR-DeepInsight enables more precise representation learning for T cell transcriptome information, better identification of general features across samples and removal of batch effects, and model transfer to new dataset. Meanwhile, BERT based deep-learning framework in TCR-DeepInsight obtained accurate TCR representation which allows clustering for CDR3α/β with variable length and amino acid mismatches. The BERT model alone in TCR-DeepInsight could also be applied to unsupervised clustering of TCRα/β or query similar TCRα/β with known epitope specificity when single-cell transcriptome information was not available. Compared with previous TCR clustering methods, our TCR-DeepInsight could integrate large-scale, multimodal, and heterogeneous single-cell transcriptome and full-length paired TCRα/β chains information all at once with relatively high scalability. Most importantly, with the reference dataset collected in this study, the scoring systems embedded in TCR-DeepInsight could overlay additional biological intuitions for TCR clusters identified. Along with additional information from HLA genotype and clonal expansion levels, our TCR-DeepInsight would greatly enhance the biological meaningful information that could be extracted from single-cell immune profiling dataset, and thus shed light on the understanding of T cell biology as well as future precise immunotherapy.

### Limitations of the study

Our TCRα/β reference atlas, which includes data from 10^6^ T cells derived from 291 individuals, provides a valuable resource for understanding TCR diversity and recurrence, although this number is just a fraction of the theoretical estimation of TCR diversity that could be as high as 10^20^. Increasing the size of the dataset in the future may significantly enhance our understanding, particularly in terms of TCR recurrence. Furthermore, our current reference atlas focuses primarily on human T cells, and incorporating data from mouse models may further advance our understanding of TCR biology. Meanwhile, our TCR-DeepInsight models currently concatenate GEX and TCR embedding with a default weight hyperparameter, but do not consider their potential underlying interaction. Developing an end-to-end model in the future that better integrates GEX and TCR representation could allow for adaptive learning of model hyperparameters. Additionally, incorporating TCR structural information as model input may further enhance model performance for downstream tasks.

## METHODS

### Integrating large-scale immune profiling datasets

We have previously described a preprocessing pipeline of single-cell immune profiling datasets with linked single-cell RNA seq library (GEX library) and single-cell TCR-seq library (TCR library).^32^ In brief, cellranger (version 6.1.2) was used to obtain the gene expression count matrix (by cellranger count command) and TCR contig annotations (by cellranger vdj command) using raw sequencing reads as input. We include high-confidence T cells (hcT cells) defined as T cells passing filtering criteria on the number of captured genes and the percentage of mitochondrial genes-derived counts, and with full-length TCR sequences with VJ segment annotation and CDR3 sequence in both α and β chains from our previous collection. We further expanded our datasets to the level of millions of high-confidence T cells by appending 249 biological samples. We annotated each sample with the individual ID, disease condition, and tissue origin. We used arcasHLA (version 0.5.0, IMGT reference version 3.46.0) to extract the HLA genotype of each individual using the aligned BAM files of the GEX library. After merging biological samples from the same individual, we annotated each individual with genotypes of HLA-A, HLA-B, HLA-C, HLA-DPB1, HLA-DRB1, and HLA-DQB1.

We integrated the transcriptome features of the hcT cells including both CD4 and CD8 T cells by learning the batch-corrected latent embedding using VAE (See VAE part of the methods). We annotated the hcT cells with CD4 or CD8 by gene count of *CD4*, *CD8A*, and *CD8B*, followed by the nearest neighbour classifier (n_neighbors=13) to categorise cells with double-positive or double-negative *CD4* or *CD8* expression into CD4 or CD8 T cells. The CD4 T cells were further categorized into naïve CD4 T cells, conventional memory CD4 T cells, and T regulatory cells by key marker genes including *SELL* (Selenoprotein L), *TCF7* (Transcription Factor 7), and *FOXP3* (Forkhead Box P3). The CD8 T cells were further divided into naïve CD8 T cells and memory or effector CD8 T cells based on the expression of *SELL* and *TCF7.* We merged the T cells with exactly the same α and β chains defined by TRAV-CDR3α-TRAJ and TRBV-CDR3β-TRBJ in each individual and obtained 635,393 unique TCRα/β sequences. Each unique TCRα/β was annotated with T cell subtypes by their deriving cells. The transcriptome feature of each T cell was visualized by projecting the learned latent embedding into 2-dimensional space using the UMAP algorithm (**McInnes et al., 2018**).^62^

### Identifying enriched VJ segment in T cell subtype

We calculated the odds ratio and *p*-value using Fisher’s exact test to discover T cell subtype-specific VJ joining in α and β chains and TRAV-TRBV combination. The odds ratio of a given VJ combination (C_VJ_) in a set of T cells of same subtype (T_type_) against all other T cells (T_others_) is given by:

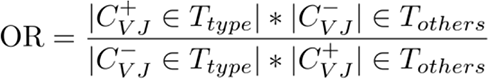

The calculation of *p*-value followed by multiple testing correction by Bonferroni method was achieved by the SciPy library (www.scipy.org). We reported VJ combinations preferentially selected by certain T cell type by thresholding the adjusted *p*-value < 0.05, odds ratio > 2, and the number supporting cells greater than 400. The selected cell type enriched VJ combination are visualized by Sankey plot (https://d3-graph-gallery.com/sankey.html) using number of cells of edges of each VJ combination and density plot in the 2-dimensional embedding of the transcriptome feature of T cells using the *kdeplot* function from the seaborn (https://seaborn.pydata.org/) library.

### Analysis of amino acid usage in the middle region of CDR3 sequence

The middle region of the CDR3 sequence in α or β chain (CDR3α-mr and CDR3β- mr) was defined as amino acid encoded by random nucleotide insertions between the V and J segments. After removing the amino acids derived from V and J segment of the CDR3 sequence, the remaining amino acids were annotated with the CDR3-mr sequence. The percentage of amino acid usage grouped by their chemical properties was calculated for each TCR, and then averaged across different T cell subtypes (CD8 T, CD4 T_conv_ and CD4 T_reg_). The significance of the difference in the percentage between different T cell subtypes was calculated by paired two-tailed Student’s t-test.

### Building reference of paired TCRa/β with CDR3 amino acid sequence and **epitope specificity**

We collected datasets with experimentally validated TCR-pMHC binding pairs. We included dataset using pMHC-tetramer cell sorting together with single-cell paired TCRα/β amplification.^22^ Paired TCRα/β with CDR3 amino acid sequences were obtained from curated databases including McPAS-TCR^45^, VDJdb^63,64^ and TCRdb^65^. We also include recently released datasets using combined DNA-barcoded pMHC tetramer/dextramer and single-cell RNA-sequencing (https://www.10xgenomics.com/cn/resources/datasets/cd-8-plus-t-cells-of-healthy-donor-1-1-standard-3-0-2, https://www.10xgenomics.com/cn/resources/datasets/cd-8-plus-t-cells-of-healthy-donor-2-1-standard-3-0-2, https://www.10xgenomics.com/cn/resources/datasets/cd-8-plus-t-cells-of-healthy-donor-3-1-standard-3-0-2, https://www.10xgenomics.com/cn/resources/datasets/cd-8-plus-t-cells-of-healthy-donor-4-1-standard-3-0-2).^46,47^ We merged the curated datasets described in deepTCR.^27^ After removing repetitive data records and data curation, we obtained 28,641 TCR-pMHC pairs with paired CDR3α and CDR3β amino acid sequences and 1037 peptide epitopes.

### Definition and analysis of public TCRs

Public TCRs were defined independently by either the single α or β chain or paired α/β chains. Public TCRα/βs were TCRs that occur in at least two individuals who share the exact same TRBV, CDR3β amino acid sequence, TRBJ, TRAV, CDR3α amino acid sequence, and TRAJ, while public TCRα and TCRβ only requires exact same VJ gene and CDR3 amino acid sequence of single α or β chain. The V and J segment call adopted the annotation from the output of cellranger. Disease-specific public TCRs were defined as TCRs occurring in at least two individuals with the same disease type.

We calculated the generation probability of α and β chains for each public TCRs and non-public TCRs by OLGA (version 1.2.4^44^). The generation probability models for the α and β chains initialized by *GenerationProbabilityVJ* and *GenerationProbabilityVDJ* function accepting both CDR3 amino acid sequence and VJ gene as input. The generation probability was followed by negative log-transformation to get the logP_gen_ of each TCR. The significance of the difference in logP_gen_ between public and non-public TCRs was calculated by paired two-tailed Student’s t-test.

### Analysis of a and β chain coherence of TCRs

The probability of independent TCRs (non-clonally expanded) of the same TRBV and CDR3β amino acid sequence that used the same TRAV gene was defined as the TRAV coherence of TCRs. In contrast, for independent TCRs of the same TRAV and CDR3α amino acid sequence that use the same TRBV gene were defined as the TRBV coherence. The TRAV and TRBV coherence were calculated separately for the CD4 T_reg_, CD4 T_n_, CD8 T_n_, CD4 T_conv_, and CD8 T memory cells separately. The definition of TRAV and TRBV coherence were also expanded to the case where the mismatch of CDR3 amino acid sequence was measured by the Levenshtein distance. The extended TRAV coherence was measured by grouping TCRβ chains with the same TRBV and mismatched CDR3β and calculating the percentage of the same TRAV in the corresponding TCRα chains. The α and β chain coherence were also measured by CDR3 sequence similarity, defined as the mean Levenshtein distance in the CDR3 sequence of one chain given that the V and CDR3 sequence was the same in the other chain.

### Training VAE model for universal representation of transcriptome features

We implemented a VAE model to capture the variation in gene expression of the 1,017,877 T cells. The probabilistic variational autoencoder accepted raw count matrices as input. The encoder received the gene expression raw count matrix and outputs a scalar mean (µ) and variance (σ^2^), which reparameterized and approximated the latent variational distribution (z). The latent distribution was regularized using the Kullback-Leibler divergence with a standard normal distribution prior (N(0,1)). To reflect biological variance and eliminate batch effects, the decoder combined the categorical encoding of the sample name, total library size by the sum of gene expression counts, and the latent embedding of the encoder output. The decoder outputed the mean, variance, and the gene-dropout probability to parameterize the zero-inflated negative-binomial distribution (ZINB) and models the raw count data.

We removed the mitochondrial genes (gene name starts with ‘MT-’), ribosomal genes (gene name starts with ‘RPS’ or ‘RPL’) and heat-shock proteins (gene name starts with ‘HSP-’). The top 6000 highly variable genes were selected using scanpy’s `pp.highly_variable_genes(flavor=’seurat_v3’, batch_key=’sample_name’, n_top_genes=6000)` before VAE integration. The VAE model was trained using the AdamW optimizer with a learning rate of 0.0001 and a batch size of 128.

### Training BERT model for universal representation of TCRa/β sequences

We adopted the BERT model originated in the field of natural language processing to obtain a scalable representation of the TCR sequence. The transformer-based models were shown to outperform conventional encoding model including MLP, convolutional networks, and recurrent models, using attention mechanism to capture inter-relationship between tokens. The BERT model was implemented in Python using the PyTorch framework and Hugginface’s Transformer libraries, with hyperparameters adapted from the default settings (hidden dimensionality = 768, intermediate size = 1536, number of attention heads = 12, number of hidden layers = 12). We implemented a tokenizer combining both CDR3α and CDR3β amino acid sequence, adding a classification token (CLS) ahead of CDR3α, gap tokens (GAP) between the two CDR3 sequence, and padding tokens (PAD) after CDR3β to align CDR3α and CDR3β sequence to the same encoding length. During pre-training, we retrieved CDR3α and CDR3β amino acid sequence from both our dataset and the curated datasets with pMHC binding information. We filtered TCRs with abnormally long CDR3α or CDR3β amino acid sequence (>36 aa), or with residues not included in the set of 20 standard amino acids, and finally retained 663,379 TCR sequence for pretraining BERT. We trained the model to predict the masked amino acids after pretraining BERT using a masked language model (MLM) objective in which we randomly hide 15% of amino acids in either CDR3α or CDR3β. We also masked either TRAV or TRAJ gene and require the model to impute the V gene usage given another chain. The MLM objective was to help the model to learn the underlying pairing pattern of TRAV/CDR3α and TRBV/CDR3β. The BERT model was pre-trained using the AdamW optimizer (a first-order stochastic optimizer) with a learning rate of 0.00001 and a batch size of 32.

The embeddings of the token from the TRAV, TRBV, CDR3β and TRBJ was used to represent the whole TCRα/β after mean pooling operation. The 768-dimensional embedding BERT output could be projected into 64-dimensional space via PCA by using the implementation of scikit-learn. The 64-dimensional representation of the TCR was then called the TCR embedding. We projected the TCR embeddings of the 634,252 TCRs to the 2-dimensional UMAP space via the UMAP algorithm using the implementation of umap-learn^62^ and visualized the embedding performance of the pretrained model.

### Unsupervised TCRa/β clustering with disease-association and shared transcriptome state

We aggregated the GEX embedding from the VAE model for each unique TCRα/β. We concatenated the TCR embeddings and the aggregated GEX embeddings and obtained a TCR-GEX joint representation of TCRα/β pairs. We adopted faiss-gpu, a computational framework for rapid similarity search optimized and accelerated by GPU^66^, for indexed *k*-nearest neighbor search of TCR-GEX joint representation based on Euclidean distance.

We use the following strategy to cluster disease-specific TCRs with transcriptome similarity. First, for each TCRα/β in our datasets, we use the TCRα/β as an anchor and find the *k*-nearest neighbor (k=40 in default). We select the top n TCRα/β pairs ranked by the Euclidean distance with the same disease, and group the anchor TCRα/β and the neighbor TCRα/βs as a potential disease-associated cluster. In the following neighbor search, we removed the possibility of the neighbor TCRα/β to prevent repeated neighbor TCRα/βs clusters.

We defined the distance between TCRs in our TCR-GEX joint representation space as the Euclidean distance between the representation of TCRs.

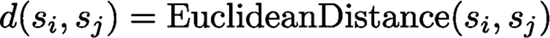

We defined a similarity score to measure the sequence similarity of TCRα/β within a disease-associated cluster. The similarity score was defined as the mean Euclidean distance in the TCR-GEX joint representation space between all neighboring TCRα/β (*s_i_*) and the anchor TCR (*s_anchor_*):

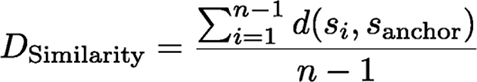

where *n* is the number of TCRs in the cluster and *s_anchor_* is the anchor TCR in the cluster.

We also calculated the mean pairwise Levenshtein distance of either CDR3α or CDR3β within a disease-associated cluster by:

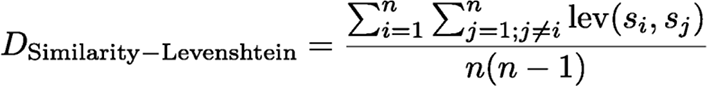

where *lev* is the Levenshtein distance of two sequences.

We defined a disease-specificity score to measure the distinctiveness of TCRα/β in a disease-associated cluster. The score was computed based on the disparity between the mean Euclidean distance in the TCR-GEX joint representation space. Specifically, the score was calculated by comparing (1) the mean Euclidean distance between the anchor TCRα/β and all neighboring TCRα/β in the cluster, and (2) the mean Euclidean distance between the top-ranked TCRs that are most similar to the anchor TCRα/β, but derived from different diseases. The latter was determined with an equal number of neighbors of the anchor TCRα/β.

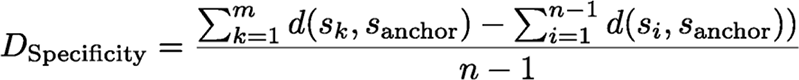

where *m* equals to *n-1*, which is the number of neighbors of the anchor TCRα/β. By using this score, we could better evaluate the uniqueness of TCRα/β in a disease-associated cluster. The proposed score enables a more accurate evaluation of the uniqueness of TCRα/β in a disease-associated cluster. Furthermore, the study also calculated the difference between the mean Levenshtein distance to the anchor TCR of either the CDR3α or the CDR3β from the disease-associated cluster and the top-ranked TCRs.

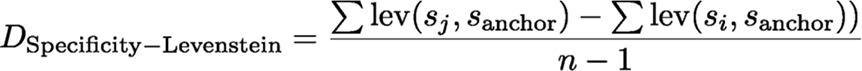

The correlation between the TCR similarity score and disease specificity and the Levenshtein distance is visualized by nonparametric loess regression.

### Selecting top-ranked disease-associated TCRa/β clusters with shared transcriptome features

The selection of disease-associated TCRα/β clusters involved the use of previously established similarity and disease-specificity scores. Specifically, TCRα/β clusters with a combined score (square of similarity score plus square of disease-specificity score) less than 300^2^ are selected for further analysis. Additional criteria are applied based on the number of unique individuals and unique TCRα/β in a TCRα/β cluster, which varied depending on the disease type. For disease types with a relatively larger number of individuals, such as solid tumors, COVID-19, and colitis, TCRα/β clusters with more than 2 unique individuals and more than 3 unique TCRα/βs were selected. For diseases with smaller individual numbers and derived from distinct studies, such as ankylosing spondylitis, TCRα/β clusters with more than 1 unique individual and more than 2 unique TCRα/βs were selected. To augment the selected TCRα/β, additional information was included, such as the common HLA genotypes of the individuals from which they were derived and the level of clonal expansion. These information served to provide an additional layer of potentially disease-related features to the TCRα/β clusters.

### Querying TCRs with known epitope specificity in the TCR atlas

We adopted the paired TCRα/β with VJ gene usage information and CDR3 amino acid sequences with known epitope specificity to search for similar TCRs in the reference atlas. We used the pre-trained BERT model to transfer TCR embeddings from the reference atlas to the query TCRs, which were then subjected to PCA transformation using the same weights as the reference atlas. We concatenated the resulting TCR embeddings from the query and reference atlas and used the query TCRs as anchors in a kNN search. We ranked the TCRs in the reference atlas by their distance to the anchor TCR for each query TCR, with distance cutoffs of 100 and 186 used for TCRs in **Figures7B** and **7C**, respectively.

### Query of new paired scRNA-seq and scTCR-seq samples against existing reference

We implemented an architecture surgery strategy to apply transfer learning in the VAE model. Specifically, during data transfer, the weights of encoder and decoder were kept the same as the VAE model before transfer learning, and the original set of batch indexes would be extended to allow extended datasets. We adopted a recently published scRNA-seq and scTCR-seq dataset including 59,417 T cells derived from 10 individuals with gastric cancer as example.^52^ As the data processing pipeline was the same as our reference atlas, we kept the same 6000 HVGs as in our previous analysis to perform transfer learning. We used a k-Nearest classifier (`n_neighbors=13`) to transfer the original cell type labels including CD8 T_n_, CD8 T, CD4 T_n_, CD4 T_conv_ and CD4 T_reg_, and MAIT to the query dataset.

We again aggregated the transferred GEX embedding by unique TCRs and concatenated it to the TCR embedding from the BERT model followed by PCA transformation with the same weight as the reference atlas. Categorizing Gastric Cancer associated TCRα/β clusters follows the same strategy described above, and TCRα/β clusters with more than 1 unique individual and more than 2 unique TCRα/βs were selected.

### Code and dataset availability

The reference atlas datasets utilized for analysis can be accessed on Figshare (excluding the two controlled access data) at https://figshare.com/collections/_/6471289. The code for TCR-DeepInsight and data analysis is available on GitHub at https://github.com/WanluLiuLab/TCR-DeepInsight, along with a Jupyter Notebook at https://huarc.net/notebook.

## ACKNOWLEDGEMENTS

We thank all the researchers who generated the single-cell immune profiling datasets used in this study. We thank Dr. Hussein A. Abbas for kindly sharing the raw data from AML patients generated in their previous publication (EGAD00001007672/EGAD0001007674).^67^ We thank Dr. Zheng Wang for kindly sharing the raw data from Kawasaki disease generated in their previous publication.^18^ We thank Dr. Hongbo Hu from Sichuan University, Dr. Feng Wang from Shanghai Jiaotong University, Dr. Chaochen Wang from Zhejiang University and all lab members from the Liu lab at ZJU-UoE Institute for their helpful discussion. We would also like to thank the technical support provided by the Core Facilities, especially the ZJE server of ZJU-UoE Institute. This work is supported by National Natural Science Foundation of China 31930038 (to L.L.), 32100718 (to L.L.), U21A20199 (to L.L.), 32170551 (to W.L.), the Fundamental Research Funds for the Central Universities 226-2022-00134 (to W.L.), the Tencent AI Lab Rhino Bird Research Funding RBFR2022015 (to W.L.) and Pre-research Projects of Innovation Center of Yangtze River Delta, Zhejiang University (to L.W., W.L., L.L.).

## AUTHOR CONTRIBUTIONS

W.L., L.L., J.Y., L.W., Z.X., and L.W., conceived the study and designed experiments. Z.W., W.L., and L.L., wrote the manuscript. Z.X., L.W., Y.B., D.S., Y.G., and P.C. processed the data. Z.X., L.W., R.T., performed the bioinformatics analysis. Z.X., Z.L., Y.Z., B.H., J.Y. and W.L., conceived the deep-learning framework. Z.W. implemented the deep-learning framework. All authors contributed to the review and corrections of the manuscripts.

## DECLARATION OF INTERESTS

The authors declare no competing financial interests.

---

**Figure S1.**
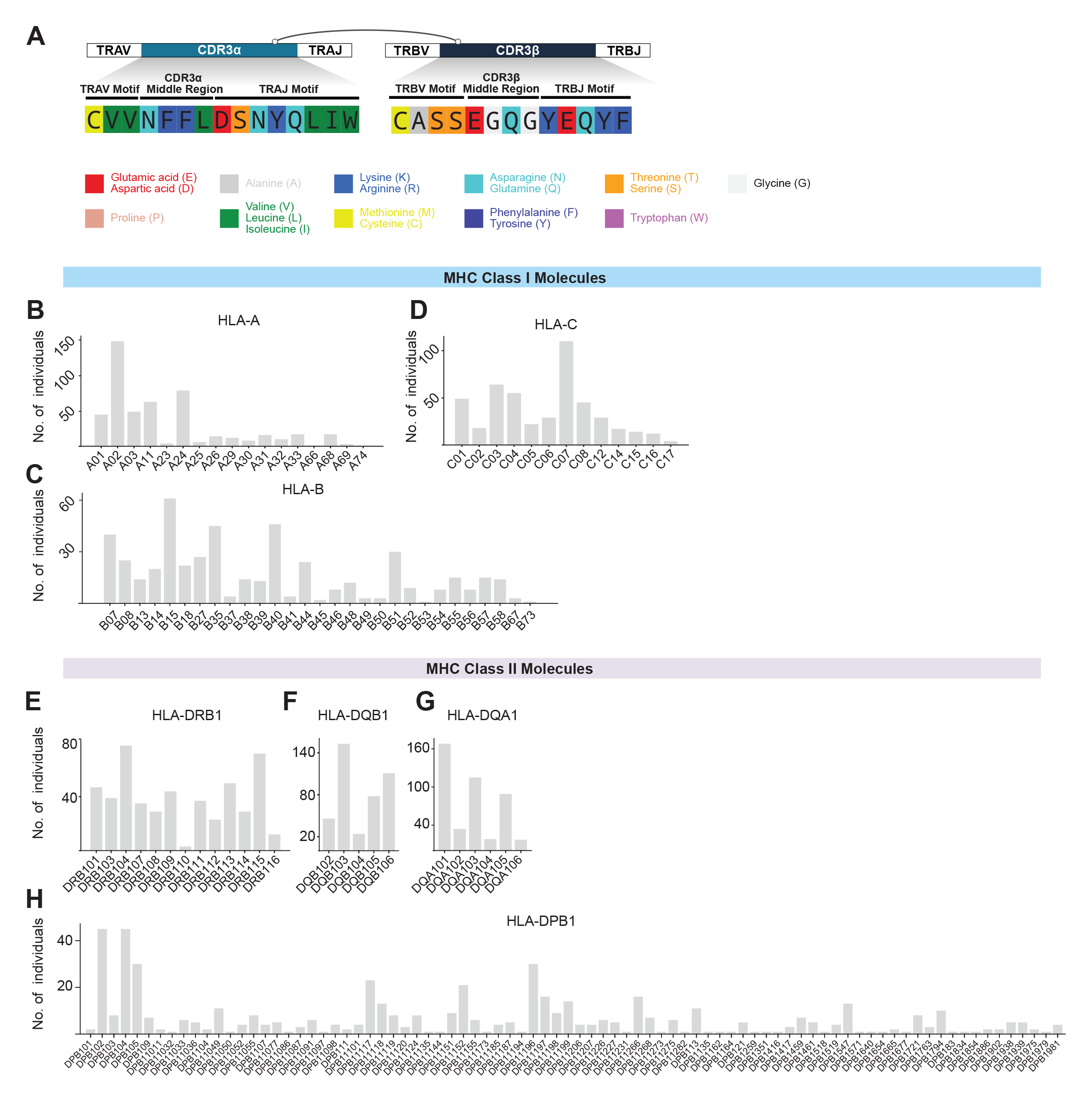
HLA Information in the integrated Paired Transcriptome and Full-Length TCR Sequences Dataset, related to Figure 1. **(A)** Schematic illustration for V gene motif, middle region, and J gene motif definition of the CDR3 region. Amino acids in this study were colored by RasMol amino acid color scheme based on Robert Fletterick’s “Shapely models”. **(B-H)** Distribution of number of individuals of called HLA alleles of MHC I HLA-A (**B**), HLA-B (**C**), HLA-C (**D**) and MHC II HLA-DRB1 (**A**), HLA-DQB1 (**F**), HLA-DQA1 (**G**), HLA-DPB1 (**H**) in the integrated datasets.

**Figure S2.**
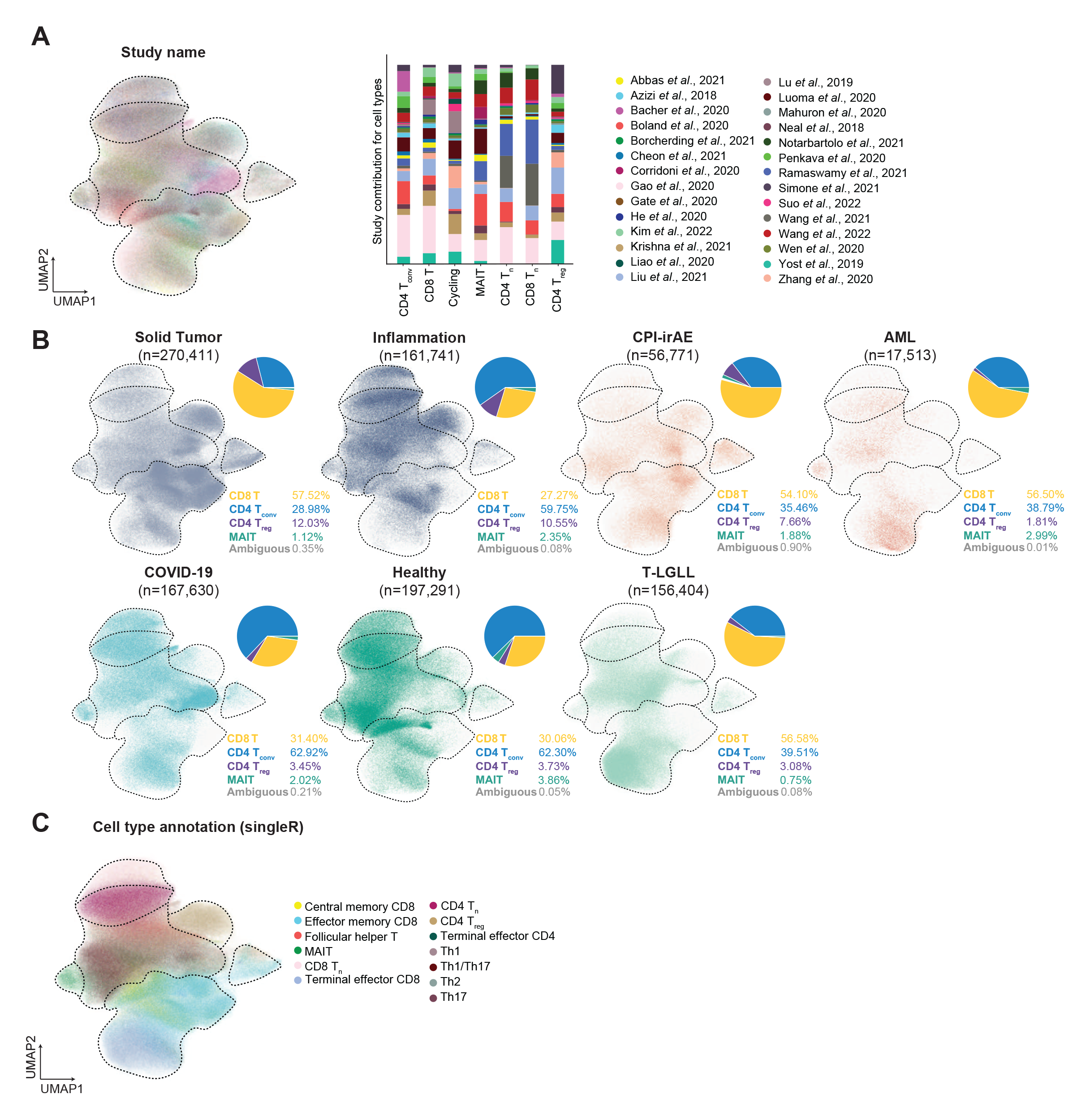
Additional information on the integrated transcriptome features of over one-million T cells, related to Figure 1. **(A)** The UMAP projection of the integrated million-level T cells colored by study names (left panel), and bar plot for the distribution of different studies corresponding to major T cell types including CD4 T_conv_, CD4 T_reg_, CD4 T_n_, CD8 T_n_, and MAIT cells (right panel). **(B)** Distribution of cells from different disease states including solid tumors, inflammation, CPI-irAE (checkpoint inhibitor associated immune-related adverse events), AML (Acute Myeloid Leukemia), COVID-19, Healthy, and T-LGLL (T cell large granular lymphocytic leukemia) in the integrated UMAP and pie chart for the composition of T cell states in each disease conditions. **(C)** Cell type annotation by singleR as previously used in *Wu et al.*, 2021. Th1: T helper 1 cell; Th17: T helper 17 cell; Th2: T helper 2 cell.

**Figure S3.**
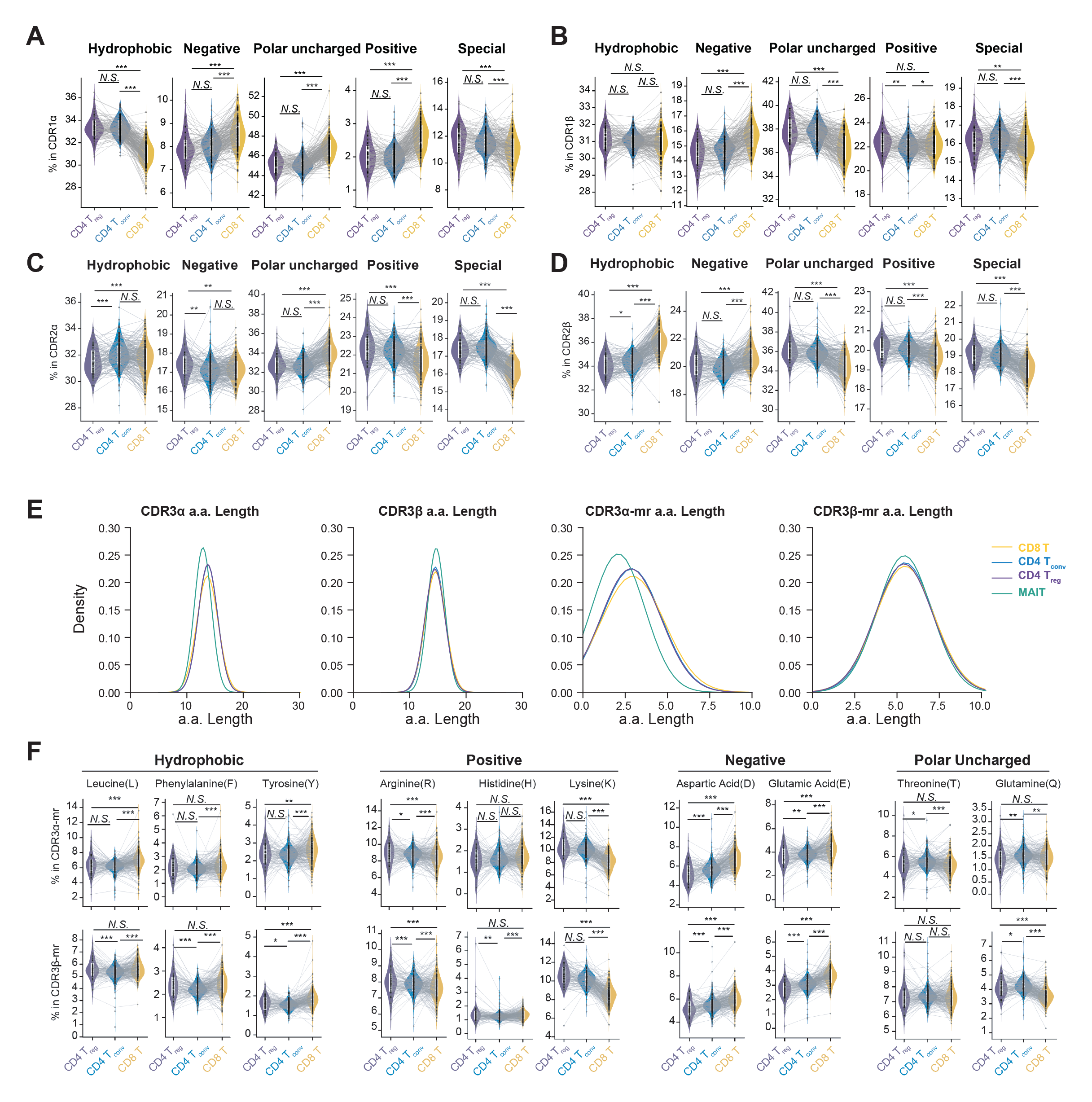
Amino acid usage and length distribution of CDR3 region of different T cell types, related to Figure 2. **(A-D)** Violin plot indicating the preference for amino acids with different physicochemical properties in CDR1α (**A**), CDR2α (**B**), CDR1β (**C**), CDR2β (**D**) regions in CD8, CD4 T_conv_, and CD4 T_reg_ cells (*** *p-value* < 0.001, ** *p-value* < 0.01, * *p-value* < 0.05, paired *t*-test; N.S.= not significant). **(E)** Distribution of the length of CDR3 and CDR3 amino acids (a.a.) mr in CD8, CD4 T_conv_, and CD4 T_reg_ cells. **(F)** Violin plot indicating the preference for amino acids with selected amino acids grouped by physicochemical properties in CDR3α-mr (upper panel) and CDR3β-mr (lower panel) in CD8, CD4 T_conv_, and CD4 T_reg_ cells (*** *p-value* < 0.001, ** *p-value* < 0.01, * *p-value* < 0.05, paired *t*-test; N.S.= not significant). Each grey dot represent one individual, lines connecting grey dots indicate amino acid usage in different cell types within one individual. White boxes within violin plot represent the distribution from 25 percentile to 75 percentile.

**Figure S4.**
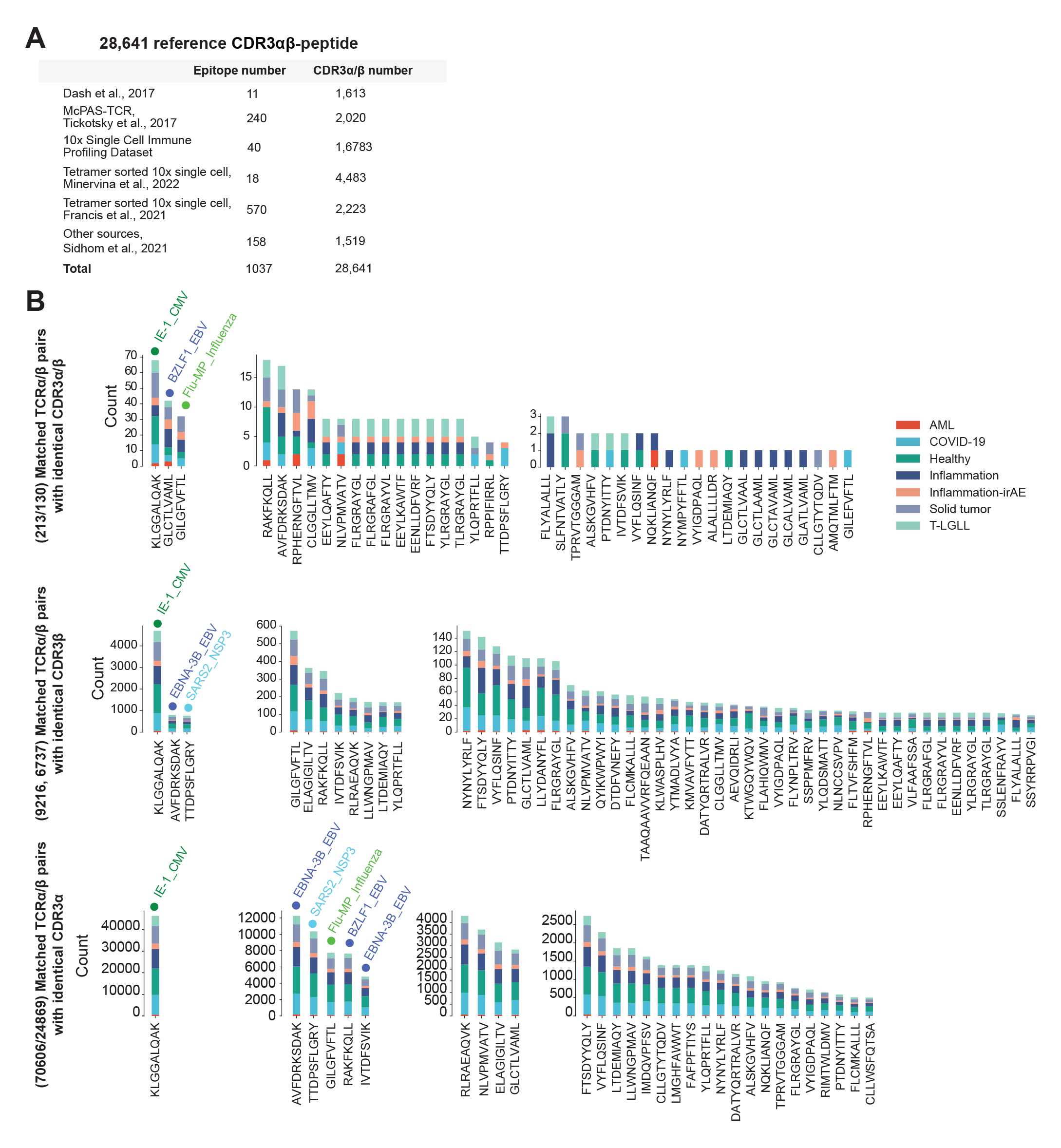
Public dataset collected for TCRα/β pairs with known epitope specificity, related to Figure 4 and 7. **(A)** Collection of reference TCR-pMHC pairs from various resources containing CDR3α, CDR3β amino acid sequence and the antigen peptide. **(B)** Number of matched TCRα/β pairs with identical CDR3α (upper panel), CDR3β (middle panel), or both CDR3α and CDR3β (lower panel) amino acid sequence with the reference TCR-pMHC information.

**Figure S5.**
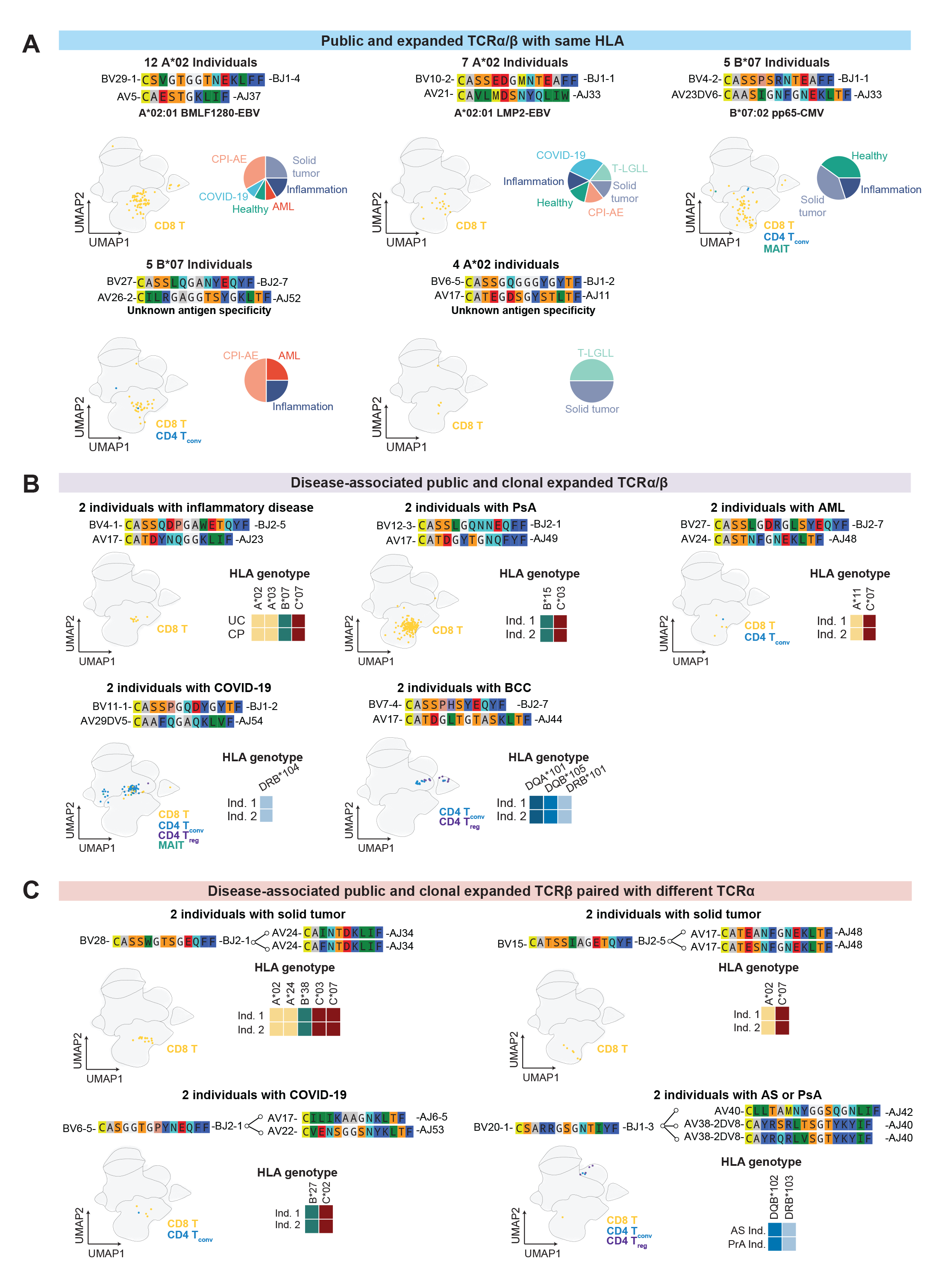
Visualization for public TCRs with their corresponding transcriptome states with at least one common HLA allele, related to Figure 4. **(A-C)** Representative public and expanded TCRα/β pairs (**A**), disease-associated public and clonal expanded TCRα/β pairs **(B)** and disease-associated public and clonal expanded TCRβ paired with different TCRα (**C**).

**Figure S6.**
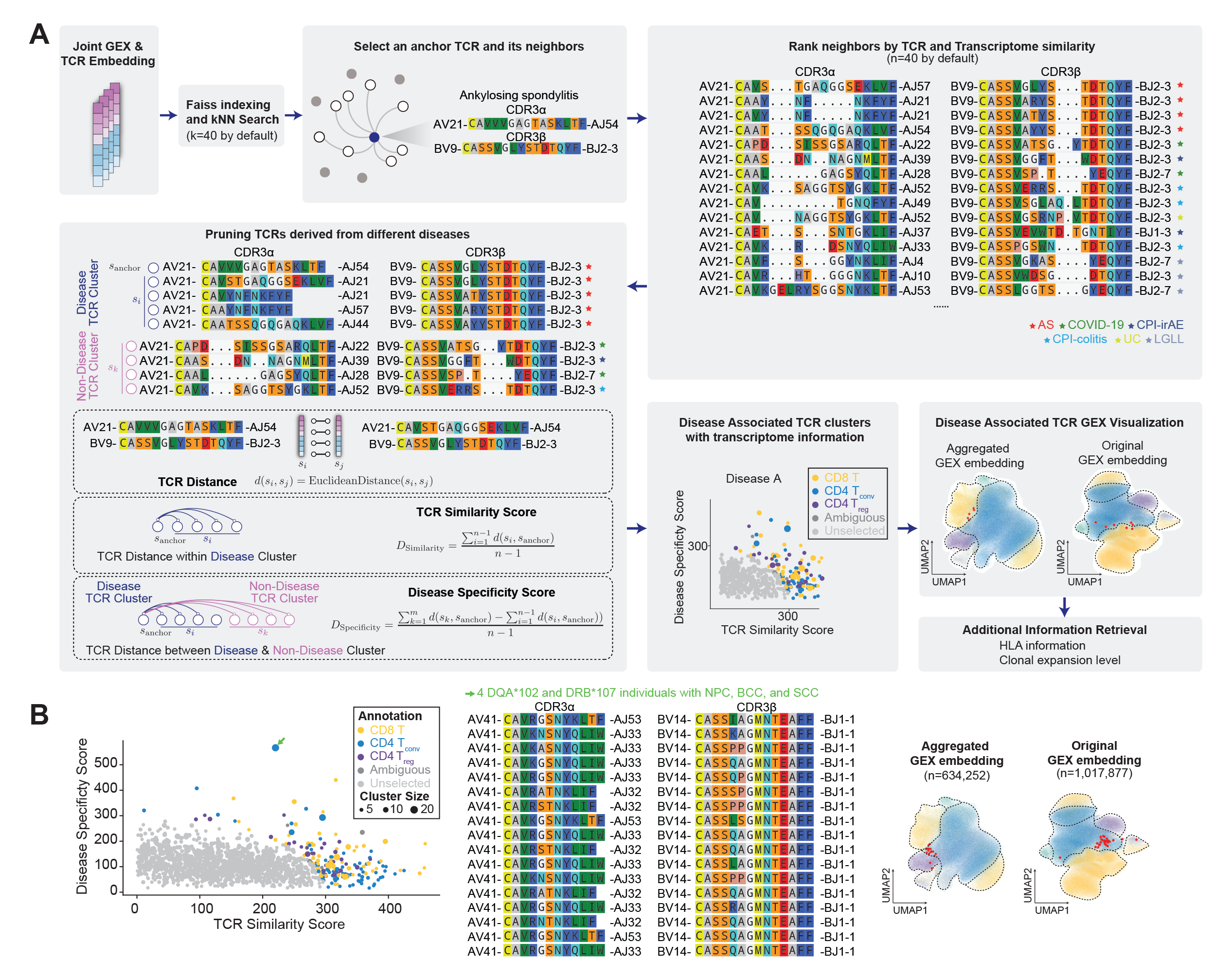
Model details for TCR-DeepInsight and representative CD4 T_conv_ dTCR cluster for solid tumor, relate to Figure 5 and 6. **(A)** Schematic illustration for the identification of dTCR clusters with TCR-DeepInsight. Briefly, joint GEX and TCR embed-dign were faiss indexed for kNN search. Then anchor TCR and its neighbors were identified and ranked by their TCR and transcriptome similarity with anchor TCR. TCRs derived from different disease were pruned to group TCRs into disease TCR cluster and non-disease TCR cluster.TCR-similarity score and disease-specificity score were calculated based on TCR distance, dTCR clusters were then indentified and visualized on GEX embedding, and additional HLA information and clonal expansion level could be retrieved. **(B)** Representative CD4 T_conv_ disease-associated TCR cluster identified in solid tumor. Scatter plot of TCR similarity score, disease specificity score, and cell type annotation for dTCR clusters were shown (left panel), VJ gene usage and CDRα/β amino acid sequences were for selected dTCR clusters were shown (middle panel), and their transcriptome featurs on original and aggregated GEX embedding were visualized (right panel). Green arrows on scatter plot indicate dTCR clusters selected for visualization with no known antigen specificity. Red stars on aggregated and original GEX UMAP indiate the cells harboring these dTCRs.

## Notes

### Competing Interest Statement

The authors have declared no competing interest.

### Summary of Updates

Add Figure S4

