## Supplementary Tables for "Disease associated human TCR characterization by deep-learning framework TCR-DeepInsight": SupplementaryTable1.docx

| **Study name** | **Number of T cells** | **Disease** |
| --- | --- | --- |
| Abbas et al., 2021 | 20302 | Acute Myeloid Leukimia |
| Azizi et al., 2018 | 20851 | Breast Cancer |
| Bacher et al., 2020 | 39950 | COVID-19 |
| Boland et al., 2020 | 84076 | Ulcerative colitis and Healthy controls |
| Borcherding et al., 2021 | 8494 | Clear cell renal cell carcinoma |
| Cheon et al., 2021 | 16271 | COVID-19 |
| Corridoni et al., 2020 | 6915 | Ulcerative colitis and Healthy controls |
| Gao et al., 2020 | 210216 | Large granular lymphocyte leukemia and healthy controls |
| Gate et al., 2020 | 871 | Healthy controls |
| He et al., 2020 | 34058 | Healthy |
| Kim et al., 2022 | 51343 | Checkpoint inhibitor associated arthritis |
| Krishna et al., 2021 | 3635 | Clear cell renal cell carcinoma |
| Liao et al., 2020 | 76477 | COVID-19 |
| Liu et al., 2021 | 34767 | Nasopharyngeal carcinoma |
| Lu et al., 2019 | 57220 | Metastatic colorectal cancer |
| Luoma et al., 2020 | 4201 | Checkpoint inhibitor associated colitis and no colitis |
| Mahuron et al., 2020 | 4315 | Melanoma |
| Neal et al., 2018 | 31568 | Clear cell renal cell carcinoma |
| Notarbartolo et al., 2021 | 36558 | COVID-19 and healthy controls |
| Penkava et al., 2020 | 56281 | Psoriatic arthritis |
| Ramaswamy et al., 2021 | 29207 | SARS-CoV-2-associated multisystem inflammatory syndrome and healthy controls |
| Simone et al., 2021 | 4633 | Ankylosing spondylitis |
| Suo et al., 2022 | 40115 | Healthy |
| Wang et al., 2021 | 54474 | Kawasaki disease and healthy controls |
| Wang et al., 2022 | 21664 | COVID-19 and healthy controls |
| Wen et al., 2020 | 44855 | COVID-19 and healthy controls |
| Yost et al., 2019 | 24560 | Basal cell carinoma and squamous cell carcinoma |
| Zheng et al., 2020 | 20302 | Esophagus squamous cell carcinoma |

**Supplementary Table 1.** Sample collections. Related to Figure 1.
