## Supplementary Tables for "Disease associated human TCR characterization by deep-learning framework TCR-DeepInsight": SupplementaryTable2.docx

| **Cell type** | **Segment1** | **Segment2** | **Odds ratio** | ***p*-value** | **Adj. *p*-value** | **Cell type^+^**  **Segment^+^** | **Cell type^-^Segment^+^** | **Cell type^+^**  **Segment^-^** | **Cell type^-^Segment^-^** |
| --- | --- | --- | --- | --- | --- | --- | --- | --- | --- |
| CD4 | TRAV23DV6 | TRAJ34 | 2.02 | 1.45E-10 | 3.91E-06 | 366 | 93 | 411075 | 210813 |
| CD4 | TRAV23DV6 | TRAJ45 | 2.40 | 3.21E-15 | 8.84E-11 | 393 | 84 | 411075 | 210813 |
| CD4 | TRAV23DV6 | TRAJ54 | 2.20 | 1.97E-12 | 5.35E-08 | 365 | 85 | 411075 | 210813 |
| CD4 | TRAV23DV6 | TRAJ22 | 2.26 | 1.99E-16 | 5.49E-12 | 475 | 108 | 411075 | 210813 |
| CD4 | TRAV13-2 | TRAJ54 | 2.02 | 2.43E-10 | 6.53E-06 | 358 | 91 | 411075 | 210813 |
| CD4 | TRAV23DV6 | TRAJ58 | 2.35 | 6.96E-18 | 1.93E-13 | 485 | 106 | 411075 | 210813 |
| CD4 | TRAV8-4 | TRAJ39 | 2.44 | 1.38E-16 | 3.80E-12 | 419 | 88 | 411075 | 210813 |
| CD4 | TRAV23DV6 | TRAJ53 | 2.66 | 2.57E-18 | 7.12E-14 | 405 | 78 | 411075 | 210813 |
| CD4 | TRAV9-2 | TRAJ22 | 2.05 | 6.74E-18 | 1.87E-13 | 635 | 159 | 411075 | 210813 |
| CD4 | TRAV23DV6 | TRAJ37 | 2.11 | 4.82E-15 | 1.33E-10 | 494 | 120 | 411075 | 210813 |
| CD4 | TRAV2 | TRAJ37 | 2.10 | 1.24E-16 | 3.43E-12 | 560 | 137 | 411075 | 210813 |
| CD4 | TRAV13-2 | TRAJ53 | 2 | 8.07E-14 | 2.21E-09 | 507 | 130 | 411075 | 210813 |
| CD4 | TRAV2 | TRAJ31 | 2.08 | 6.88E-13 | 1.88E-08 | 430 | 106 | 411075 | 210813 |
| CD4 | TRAV23DV6 | TRBV5-1 | 2.58 | 1.14E-39 | 3.21E-35 | 968 | 193 | 411075 | 210813 |
| CD4 | TRAV9-2 | TRBV12-3 | 2.19 | 2.61E-20 | 7.27E-16 | 639 | 150 | 411075 | 210813 |
| CD4 | TRAV9-2 | TRBV30 | 2.06 | 1.79E-16 | 4.93E-12 | 574 | 143 | 411075 | 210813 |
| CD4 | TRAV16 | TRBV5-1 | 2.01 | 7.24E-16 | 2E-11 | 584 | 149 | 411075 | 210813 |
| CD4 | TRAV23DV6 | TRBV2 | 2.32 | 2.89E-16 | 7.97E-12 | 448 | 99 | 411075 | 210813 |
| CD4 | TRAV8-4 | TRBV7-2 | 2.02 | 4.53E-24 | 1.27E-19 | 911 | 232 | 411075 | 210813 |
| CD4 | TRAV23DV6 | TRBV5-4 | 2.33 | 1.56E-16 | 4.32E-12 | 450 | 99 | 411075 | 210813 |
| CD4 | TRAV2 | TRBV7-2 | 2.35 | 5.76E-17 | 1.59E-12 | 458 | 100 | 411075 | 210813 |
| CD4 | TRAV26-1 | TRBV12-3 | 2.02 | 2.19E-12 | 5.97E-08 | 441 | 112 | 411075 | 210813 |
| CD4 | TRAV8-4 | TRBV12-3 | 2.14 | 2.24E-15 | 6.18E-11 | 489 | 117 | 411075 | 210813 |
| CD4 | TRAV13-2 | TRBV12-3 | 2.07 | 1.04E-11 | 2.82E-07 | 391 | 97 | 411075 | 210813 |
| CD4 | TRAV23DV6 | TRBV20-1 | 2.31 | 7.07E-38 | 2E-33 | 1104 | 245 | 411075 | 210813 |
| CD4 | TRAV23DV6 | TRBV7-2 | 2.30 | 2.27E-19 | 6.31E-15 | 547 | 122 | 411075 | 210813 |
| CD4 | TRAV2 | TRBV2 | 2.24 | 1.40E-12 | 3.80E-08 | 362 | 83 | 411075 | 210813 |
| CD4 | TRAV23DV6 | TRBV3-1 | 2.79 | 7.49E-20 | 2.08E-15 | 413 | 76 | 411075 | 210813 |
| CD4 | TRAV23DV6 | TRBV18 | 2.48 | 2.30E-19 | 6.40E-15 | 483 | 100 | 411075 | 210813 |
| CD4 | TRBV3-1 | TRBJ1-6 | 2.02 | 3.54E-19 | 9.82E-15 | 712 | 181 | 411075 | 210813 |
| CD4 | TRBV5-4 | TRBJ1-6 | 2.07 | 6.33E-13 | 1.73E-08 | 439 | 109 | 411075 | 210813 |
| CD8 | TRAV14DV4 | TRBV19 | 2.31 | 8.43E-39 | 2.38E-34 | 434 | 686 | 134045 | 487843 |
| CD8 | TRAV14DV4 | TRBV27 | 3.51 | 1.18E-61 | 3.35E-57 | 366 | 380 | 134045 | 487843 |
| CD8 | TRAV29DV5 | TRBV27 | 2.53 | 3.90E-45 | 1.10E-40 | 427 | 616 | 134045 | 487843 |
| CD8 | TRAV12-2 | TRBV9 | 2.01 | 1.07E-27 | 3E-23 | 411 | 744 | 134045 | 487843 |
| CD8 | TRAV17 | TRBV27 | 2.67 | 3.07E-46 | 8.68E-42 | 400 | 547 | 134045 | 487843 |
| CD8 | TRAV19 | TRBV9 | 2.98 | 2.78E-54 | 7.87E-50 | 400 | 490 | 134045 | 487843 |
| CD8 | TRAV21 | TRBV27 | 3.30 | 5.99E-70 | 1.70E-65 | 450 | 497 | 134045 | 487843 |
| CD8 | TRAV12-2 | TRBV27 | 2.92 | 6.01E-47 | 1.70E-42 | 353 | 441 | 134045 | 487843 |
| CD8 | TRAV14DV4 | TRBV9 | 2.82 | 8.85E-57 | 2.51E-52 | 452 | 584 | 134045 | 487843 |
| CD8 | TRAV1-2 | TRBV9 | 4.16 | 3.68E-74 | 1.04E-69 | 368 | 323 | 134045 | 487843 |
| CD8 | TRAV12-2 | TRBV7-9 | 2 | 9.44E-24 | 2.64E-19 | 354 | 645 | 134045 | 487843 |
| CD8 | TRAV21 | TRBV7-9 | 2.24 | 9.54E-32 | 2.68E-27 | 371 | 603 | 134045 | 487843 |
| CD8 | TRAV19 | TRBV7-9 | 2.68 | 1.60E-55 | 4.53E-51 | 480 | 653 | 134045 | 487843 |
| CD8 | TRAV19 | TRBV27 | 3.61 | 3.98E-69 | 1.13E-64 | 399 | 403 | 134045 | 487843 |
| CD8 | TRAV14DV4 | TRBV7-9 | 2.73 | 4.99E-56 | 1.41E-51 | 470 | 628 | 134045 | 487843 |
| CD8 | TRAV19 | TRBV19 | 2.13 | 1.65E-30 | 4.63E-26 | 399 | 683 | 134045 | 487843 |
| CD8 | TRBV27 | TRBJ2-7 | 2.71 | 4.82E-193 | 1.37E-188 | 1685 | 2279 | 134045 | 487843 |
| CD8 | TRBV7-9 | TRBJ2-1 | 2.03 | 3.92E-96 | 1.11E-91 | 1462 | 2630 | 134045 | 487843 |
| CD8 | TRBV4-1 | TRBJ2-7 | 2.41 | 1.29E-90 | 3.67E-86 | 960 | 1458 | 134045 | 487843 |
| CD8 | TRBV27 | TRBJ1-1 | 2.20 | 5.75E-63 | 1.63E-58 | 791 | 1312 | 134045 | 487843 |
| CD8 | TRBV27 | TRBJ2-3 | 2.14 | 5.43E-59 | 1.54E-54 | 783 | 1333 | 134045 | 487843 |
| CD8 | TRBV13 | TRBJ2-7 | 3.26 | 7.72E-64 | 2.19E-59 | 417 | 467 | 134045 | 487843 |
| CD8 | TRBV7-6 | TRBJ2-7 | 2.16 | 7.80E-37 | 2.20E-32 | 472 | 798 | 134045 | 487843 |
| CD8 | TRBV27 | TRBJ2-1 | 2.90 | 3.66E-180 | 1.04E-175 | 1411 | 1780 | 134045 | 487843 |
| CD8 | TRBV9 | TRBJ2-7 | 2.05 | 1.54E-70 | 4.37E-66 | 1045 | 1863 | 134045 | 487843 |
| CD8 | TRBV4-1 | TRBJ1-1 | 2.04 | 3.47E-36 | 9.79E-32 | 530 | 949 | 134045 | 487843 |
| CD8 | TRBV7-6 | TRBJ2-1 | 2.07 | 5.68E-30 | 1.60E-25 | 419 | 739 | 134045 | 487843 |
| MAIT | TRAV1-2 | TRAJ33 | 210.63 | 1e-320 | 1e-320 | 5049 | 2280 | 11443 | 610445 |
| MAIT | TRAV1-2 | TRAJ20 | 19.89 | 1e-320 | 1e-320 | 376 | 1041 | 11443 | 610445 |
| MAIT | TRAV1-2 | TRAJ12 | 55.87 | 1e-320 | 1e-320 | 452 | 449 | 11443 | 610445 |
| MAIT | TRAV1-2 | TRBV6-4 | 197.73 | 1e-320 | 1e-320 | 1468 | 454 | 11443 | 610445 |
| MAIT | TRAV1-2 | TRBV6-2 | 58.88 | 1e-320 | 1e-320 | 573 | 546 | 11443 | 610445 |
| MAIT | TRAV1-2 | TRBV6-1 | 66.72 | 1e-320 | 1e-320 | 917 | 796 | 11443 | 610445 |
| MAIT | TRAV1-2 | TRBV20-1 | 39.66 | 1e-320 | 1e-320 | 1074 | 1590 | 11443 | 610445 |
| MAIT | TRAV1-2 | TRBV4-2 | 51.87 | 1e-320 | 1e-320 | 582 | 630 | 11443 | 610445 |
| MAIT | TRBV6-4 | TRBJ2-3 | 77.35 | 1e-320 | 1e-320 | 619 | 451 | 11443 | 610445 |
| MAIT | TRBV6-4 | TRBJ2-1 | 40.60 | 1e-320 | 1e-320 | 455 | 622 | 11443 | 610445 |

**Supplementary Table 2.** T cell receptor (TCR) VJ segment pairing in different T cell types. Related to Figure 2.
