## Supplementary Tables for "Disease associated human TCR characterization by deep-learning framework TCR-DeepInsight": SupplementaryTable3.docx

|  | **Cell type** | **Mean** | **Variance** |
| --- | --- | --- | --- |
| CDR3a | CD4 | 13.735442 | 1.715207 |
|  | CD8 | 13.65522 | 1.88235 |
|  | MAIT | 12.831251 | 1.50513 |
|  | Treg | 13.718407 | 1.711401 |
| CDR3b | CD4 | 14.533363 | 1.74841 |
|  | CD8 | 14.56619 | 1.819232 |
|  | MAIT | 14.692301 | 1.511805 |
|  | Treg | 14.462966 | 1.779967 |
| CDR3a_mr | CD4 | 2.887274 | 1.772451 |
|  | CD8 | 2.955396 | 1.885915 |
|  | MAIT | 2.086516 | 1.569067 |
|  | Treg | 2.846795 | 1.770939 |
| CDR3b_mr | CD4 | 5.476402 | 1.690104 |
|  | CD8 | 5.487635 | 1.732546 |
|  | MAIT | 5.461854 | 1.593268 |
|  | Treg | 5.424257 | 1.703939 |

**Supplementary Table 3.** Length of CDR3a and CDR3b amino acid sequence in major cell types.
