## Supplementary Tables for "Disease associated human TCR characterization by deep-learning framework TCR-DeepInsight": SupplementaryTable4.docx

| **Region** | **Amino acid property** | **Cell type 1** | **Cell type 2** | **Mean percentage for cell type 1** | **Mean percentage for cell type 2** | **p-value paired t.test** |
| --- | --- | --- | --- | --- | --- | --- |
| CDR3α_mr | Hydrophobic | Treg | CD4 | 31.0703644 | 30.7441261 | 0.1829028 |
| CDR3α_mr | Hydrophobic | CD4 | CD8 | 30.893405 | 32.7552242 | 5.56E-23 |
| CDR3α_mr | Hydrophobic | Treg | CD8 | 31.0164181 | 32.930972 | 4.98E-10 |
| CDR3α_mr | Negative | Treg | CD4 | 8.82263789 | 9.45636845 | 3.33E-06 |
| CDR3α_mr | Negative | CD4 | CD8 | 9.40488181 | 10.9759967 | 2.75E-36 |
| CDR3α_mr | Negative | Treg | CD8 | 8.83083429 | 10.8599482 | 3.25E-21 |
| CDR3α_mr | Polar uncharged | Treg | CD4 | 15.7449227 | 16.5445524 | 0.0001039 |
| CDR3α_mr | Polar uncharged | CD4 | CD8 | 16.4166951 | 15.4864872 | 4.42E-14 |
| CDR3α_mr | Polar uncharged | Treg | CD8 | 15.7146232 | 15.3723285 | 0.09196545 |
| CDR3α_mr | Positive | Treg | CD4 | 15.9427362 | 15.536972 | 0.01719705 |
| CDR3α_mr | Positive | CD4 | CD8 | 15.5850918 | 12.983142 | 1.28E-55 |
| CDR3α_mr | Positive | Treg | CD8 | 15.965633 | 12.9621548 | 1.99E-29 |
| CDR3α_mr | Special | Treg | CD4 | 22.1415432 | 22.0414173 | 0.59464263 |
| CDR3α_mr | Special | CD4 | CD8 | 22.0188172 | 21.9701847 | 0.72114431 |
| CDR3α_mr | Special | Treg | CD8 | 22.1552237 | 21.9843835 | 0.44463519 |
| CDR3β_mr | Hydrophobic | Treg | CD4 | 26.041143 | 25.0941894 | 2.50E-08 |
| CDR3β_mr | Hydrophobic | CD4 | CD8 | 25.1502127 | 26.2141017 | 2.86E-18 |
| CDR3β_mr | Hydrophobic | Treg | CD8 | 26.0125336 | 26.0441343 | 0.8755467 |
| CDR3β_mr | Negative | Treg | CD4 | 7.86545411 | 8.59299285 | 1.24E-10 |
| CDR3β_mr | Negative | CD4 | CD8 | 8.49863636 | 9.65121834 | 2.29E-31 |
| CDR3β_mr | Negative | Treg | CD8 | 7.89026795 | 9.59493145 | 6.10E-24 |
| CDR3β_mr | Polar uncharged | Treg | CD4 | 21.337658 | 22.1056885 | 2.10E-07 |
| CDR3β_mr | Polar uncharged | CD4 | CD8 | 22.0759793 | 21.4716323 | 4.01E-09 |
| CDR3β_mr | Polar uncharged | Treg | CD8 | 21.3338856 | 21.4392731 | 0.46233257 |
| CDR3β_mr | Positive | Treg | CD4 | 13.5304771 | 12.8391569 | 8.49E-09 |
| CDR3β_mr | Positive | CD4 | CD8 | 12.9657736 | 11.0730437 | 2.71E-57 |
| CDR3β_mr | Positive | Treg | CD8 | 13.5350106 | 11.2495683 | 2.51E-27 |
| CDR3β_mr | Special | Treg | CD4 | 31.0496458 | 31.2430561 | 0.27220475 |
| CDR3β_mr | Special | CD4 | CD8 | 31.1888319 | 31.4347754 | 0.02364315 |
| CDR3β_mr | Special | Treg | CD8 | 31.0477562 | 31.5045559 | 0.01586196 |

**Supplementary Table 4.** Difference in amino acid composition in the CDR3 middle region between T cell types.
